# Seasonal dynamics and drivers of microbial communities in a temperate dimictic lake: insights from metabarcoding and machine learning

**DOI:** 10.1101/2024.01.12.575328

**Authors:** Karlicki Michał, Bednarska Anna, Hałakuc Paweł, Maciszewski Kacper, Karnkowska Anna

## Abstract

Microbial communities, consisting of prokaryotes and protists, play a central role in ecological processes in aquatic environments. To understand these communities, metabarcoding provides a powerful tool to assess their taxonomic composition and to track spatio-temporal dynamics in both marine and freshwater environments. While previous research has primarily focused on marine ecosystems, it is important to study microbial communities in freshwater environments, which are characterised by high diversity and susceptibility to rapid environmental change. In temperate lakes, despite extensive research on temporal changes in physico-chemical factors and microscopic studies of plankton, there is a notable research gap regarding their eukaryotic microbial communities. Our study fills this gap by investigating the diversity and seasonal changes of prokaryotic and eukaryotic communities in Lake Roś (Poland), a representative temperate lake characterised by two mixing episodes in spring and autumn and pronounced stratification in summer. Our metabarcoding analysis revealed that both the bacterial and protist communities exhibit distinct seasonal patterns that are not necessarily shaped by dominant taxa. To decipher the drivers of the seasonal communities, we used machine learning and statistical methods and identified crucial amplicon sequence variants (ASVs) specific to each season. In addition, we identified a distinct community in the anoxic hypolimnion. We have also shown that the key factors shaping the community composition in Lake Roś are temperature, oxygen and silicon concentration. Understanding these community structures and the underlying factors is crucial in the context of climate change, which might affect mixing patterns and lead to prolonged stratification. Given the pronounced seasonal shifts observed in these communities, we can anticipate that climate change will profoundly impact the functioning of temperate dimictic lakes.

## 1 Introduction

Protists are abundant and diverse eukaryotic microorganisms in aquatic ecosystems and fulfil critical ecosystem functions (Sherr & Sherr, 2002; Worden et al., 2015; Singer et al., 2021). They play an essential role in organic matter cycling by contributing to primary production and decomposition of organic matter and constitute a link between prokaryotes and higher trophic level organisms (Azam et al., 1983; Caron, 1994; Nakano et al., 1998; Coleman & Whitman, 2005; Boenigk & Arndt, 2002; Posch et al., 2015; Šimek et al., 2020). Despite their ecological significance, the comprehensive understanding of protist diversity remains limited. Recent advancements in molecular techniques have spurred a surge in diversity studies, revealing an unexpectedly high diversity of protists across various aquatic environments, particularly in oceans e.g. (López-García et al., 2001; de Vargas et al., 2015; Lima-Mendez et al., 2015; Lovejoy et al., 2006; Worden et al., 2006; Massana et al., 2015; Seeleuthner et al., 2018; Stoeck et al., 2010; Sunagawa et al., 2020). However, studies on freshwater protist diversity remain comparatively scarce, often focusing on specific types of water bodies e.g. (Charvet et al., 2012; Taib et al., 2013; Simon et al., 2015b; Debroas et al., 2017; Cruaud et al., 2019; David et al., 2021; Metz et al., 2022).

Freshwater ecosystems are more fragmented and isolated (Dodson, 1992; Reche et al., 2005), compared to the ocean, where microbial communities are disseminated on a global scale via ocean currents (Villarino et al., 2018; Richter et al., 2022). This intrinsic lower connectivity of freshwater ecosystems hinders the dispersal of freshwater organisms and increases their genetic diversity (Manel et al., 2020; Miller, 2021). Furthermore, freshwater ecosystems’ environmental conditions are more heterogeneous and much more sensitive to external factors than those in the oceans (Simon et al., 2015b). Recent analyses across diverse habitats revealed apparent differences in the taxonomic composition of the major protistan lineages and a higher β-diversity in freshwater bodies than in the other systems (Singer et al., 2021; Xiong et al., 2021), prompting studies on freshwater ecosystems.

Within the realm of freshwater ecosystems, lakes are the most studied (Charvet et al., 2012; Lepère et al., 2016; Boenigk et al., 2018). Notably, research has predominantly concentrated on high mountain lakes (Kammerlander et al., 2016; Filker et al., 2016; Boenigk et al., 2018) and polar lakes (Daniel et al., 2016; Stoof-Leichsenring et al., 2020) due to their extreme conditions, including temperature, nutrient availability and UV radiation. Several studies have been performed on shallow eutrophic lakes (Simon et al., 2015a; Simon et al., 2015b), lakes with anoxic hypolimnion (Oikonomou et al., 2015; Lepère et al., 2016; Fermani et al., 2021) and deep lakes with oxygenated hypolimnion (Mukherjee et al., 2017). All these diverse lacustrine ecosystems consistently reveal a substantial prevalence of unclassified sequences within numerous eukaryotic lineages. Comparatively fewer molecular biodiversity surveys have been conducted in temperate lake environments (Lefranc et al., 2005; Boenigk et al., 2018; Mitsi et al., 2023). The water mixing patterns in holomictic freshwater lakes, where the water column is mixed in some seasons and remain stratified in other seasons, results in the recurring microbial communities’ assembly processes. Deep dimictic lakes undergo mixing only during the spring and autumn months, maintaining stratification throughout the summer, and winter (Kirillin & Shatwell, 2016). However, climate changes influence lakes’ mixing regimes (Adrian et al., 2009), which might profoundly impact these ecosystems by either enhancing or impeding vertical nutrient and dissolved gas fluxes (Råman Vinnå et al., 2021). Consequently, temperate lakes, characterized by their water mixing patterns, offer a valuable opportunity to study seasonal protists’ community dynamics (Lepère et al., 2010; Medinger et al., 2010; Nolte et al., 2010; Mukherjee et al., 2017).

The PEG model (Sommer et al., 1986; Sommer et al., 2012) provides the best framework for describing the seasonal succession of phytoplankton and zooplankton in aquatic ecosystems. Several studies (Kent et al., 2007; Paver et al., 2015; Woodhouse et al., 2016; Bock et al., 2018) have shown consistent temporal dynamics between eukaryotic phytoplankton and bacteria. However, prokaryotic and protist communities may show different temporal patterns over the course of the season. These differences could be due to variations in small-scale temporal patterns where prokaryotes and protists synchronise, as opposed to large-scale patterns where synchrony decreases due to changes in environmental conditions (Tammert et al., 2015; Obertegger et al., 2019).

Decades of research have unveiled the pivotal role of physical factors, such as light, temperature and turbulence in shaping of the microbial communities (Margalef, 1978; Sommer et al., 2012; Barton et al., 2015). However, seasonal succession is also governed by biological factors including organismal interactions (Drake, 1990; Dakos et al., 2009; Logares et al., 2018; Bock et al., 2020). The advent of high-throughput DNA sequencing technologies has significantly bolstered our capacity to delineate microbial diversity and discern seasonal fluctuations within aquatic environments (Bunse & Pinhassi, 2017; Giner et al., 2019; Grossart et al., 2020). Identifying these temporal patterns and determining their principal environmental drivers are essential to revealing the mechanisms governing species succession and shaping community composition. Moreover, such investigations provide valuable insights into how climate change might alter these processes (Edwards & Richardson, 2004; Siano et al., 2021; Caracciolo et al., 2022).

In this study, we conducted a metabarcoding investigation of the prokaryotic and protist communities in the typical temperate dimictic lake. We investigated small protists (size fraction 3-12 μm) and free-living bacteria (size fraction 0.2-3 μm) during the ice-free season to determine the temporal dynamics of the community composition under pronounced seasonal gradients and to identify the main drivers of the communities during the seasons. We also investigated the influence exerted by abiotic parameters, e.g., temperature, organic carbon, and nutrient availability, as well as biotic parameters on the microbial community structure.

## 2 Experimental procedures

### 2.1 Site description

Lake Roś (area: 18.08 km^2^; maximum depth: 31.8 m, mean depth: 8.1 m) is a meso/eutrophic glacial lake situated in north-eastern Poland in the area of Masurian Lake District (53°38’-53°41’ N 21°48’-21°59’ E). It is a typical temperate dimictic lake, with biannual (spring and autumn) mixing events. During the summer, Lake Roś experiences thermal stratification, leading to a pronounced vertical gradient of dissolved oxygen (DO), ranging from oxic conditions in the epilimnion to near anoxia in the hypolimnion (Dawidowicz, 1990). The lake periodically freezes during the winter months. Lake Roś has been a focal point for extensive ecological investigations throughout the 20th century, with particular attention given to macrophytes, phytoplankton, zooplankton, macroinvertebrates, and fish (e.g. Dawidowicz, 1990; Jasser, 1995; Pieczyńska et al., 1998). The lake consists of two basins connected by a relatively narrow and shallow channel (Figure 1A). The main southern basin is deeper (with a maximum depth of 31.8 m), maintains thermal stratification throughout the summer, and experiences common oxygen deficits within the hypolimnion. The second, northern basin is shallower (with a maximum depth of 9.3 m), and is predominantly covered by submerged macrophytes. This basin frequently experiences summer destratification events, leading to complete mixing of its waters.

**Figure 1.**
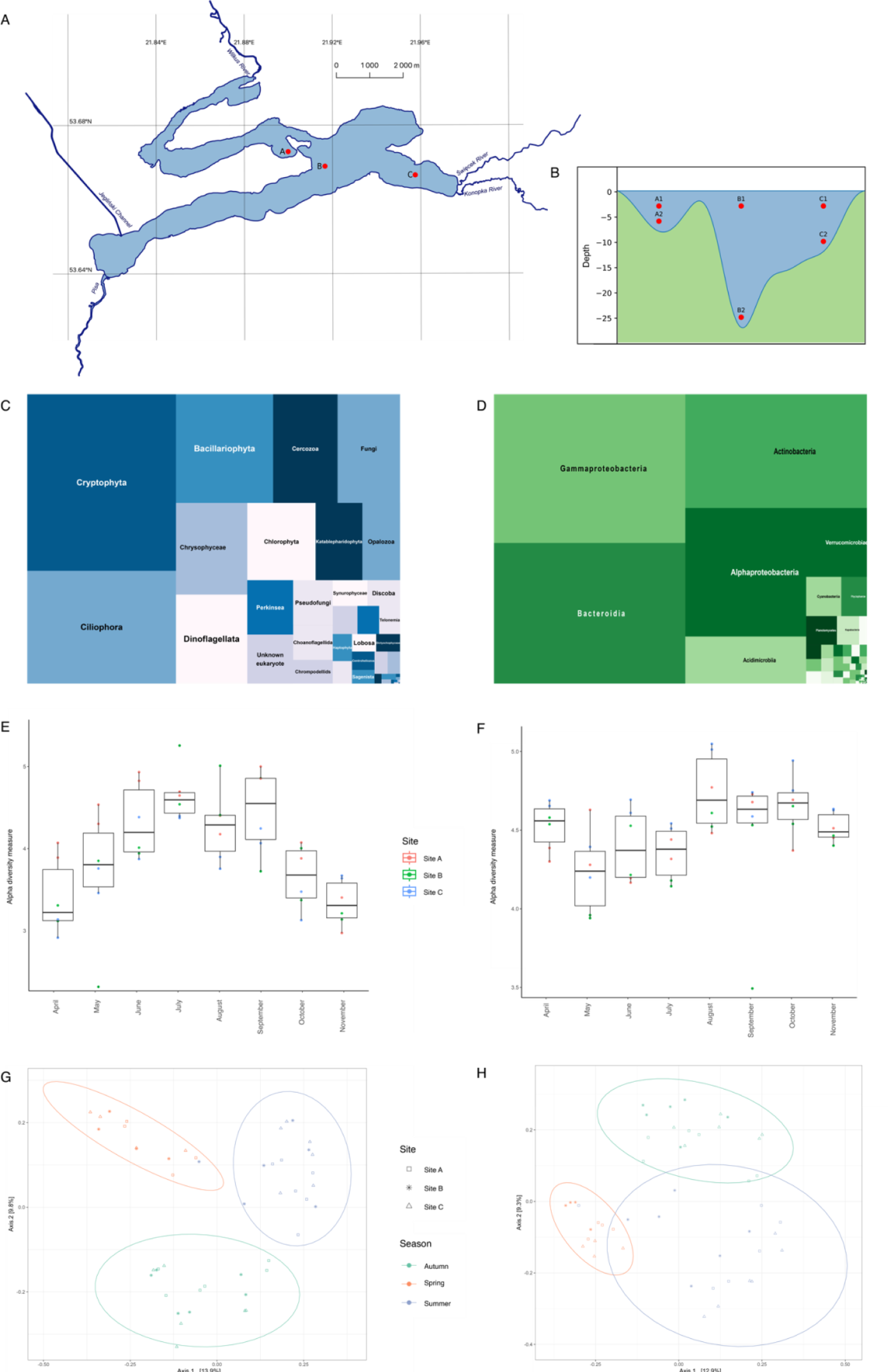
Sampling scheme and overview of the microbial community structure. Location of sampling sites A, B and C in Lake Roś (A) with the sampling depths (meters) for each sampling site (B). Treemaps represent the overall composition of relative abundances for 18S V9 rDNA at the ‘Class’ level (C) and 16S V4 rDNA at the ‘Phylum’ level (D). Boxplots illustrating Shannon alpha diversity for each month for 18S V9 rDNA (E) and 16S V4 rDNA (F) datasets, sites are colour coded. Ordination plots based on Unweighted UniFrac MDS for 18S V9 rDNA (G) and 16S V4 rDNA (H), with sampling sites marked with shapes and seasons marked with colours.

### 2.2 Sample collection

The sampling was conducted eight times during the ice-free season (from April to November) of 2019 in three sites within the lake (site A, B and C on Figure 1A), which differ in their maximum depth ranging from 8 to 27 metres (Figure 1B; Supplementary Table S1). The site A is located in the sheltered bay, in the vicinity of the deepest spot of the northern basin of the lake. Site B is located in the main basin of the lake, in the vicinity of the deepest spot of the lake. Site C is located on the periphery of the main basin, near the inflow of two rivers: Święcek and Konopka. In each site we obtained two samples of a total volume of 2L using modified Bernatowicz sampler: one from the surface layer representing euphotic zone (3 m across all sites – A1, B1, C1), while the second sample was taken from a depth of two meters above the lake bottom (A2 - 6 m, B2 25 m, C2 10 m). Samples were immediately filtered with a 150 μm plankton net to remove large particles and multicellular organisms. Further filtration has been done sequentially with minimal (up to 200 hPa above atmospheric pressure) air pressure using Nucleopore filters (Whatman, Maidstone, UK) with 12 μm, 3 μm and 0.2 μm pore size. This process continued until filter clogging was detected, allowing us to obtain size fractions of 3-12 μm and 0.2-3 μm. Filters were then securely stored in −80 °C until the DNA extraction was conducted.

Planktonic animals filtered out from the samples using 150 μm plankton net were immediately preserved with 4% formalin. Subsequently, these specimens underwent thorough examination using dissecting microscopy. Cladocerans were identified to the species level, while copepods were identified at the order level, with nauplius (larva) stages distinguished as a separate category. The densities of zooplankton taxa expressed in ind L-1 were calculated from the number of animals observed within the samples and the corresponding sample volumes.

Meteorological data for the lake area such as rainfall, and air temperature was derived from publicly available resources of the Institute of Meteorology and Water Management, Poland. *In situ* water temperature and oxygen level were measured using a YSI ProODO multiparametric probe, while the depth of the photic zone was measured using a portable light metre (LiCor LI-250A with spherical underwater PAR quantum sensor LI-193R) and the Secchi disc. *Ex situ* concentrations of biogenic (carbon, nitrogen, phosphorus) and other (iron, manganese and silicate) compounds were measured by an external company (Wessling SA) (Supplementary Table S2).

### 2.3 DNA Extraction, DNA Amplifications and Sequencing

For each sampling event and filter size (0.2 μm for prokaryotes and 3 μm for eukaryotic fraction), DNA was extracted from ¼ or ½ of the filter using the GeneMATRIX Soil DNA Purification Kit (EURx) for a total of 96 samples, according to manufacturer protocol with minor modifications. Extracted environmental DNA was quantified using NanoDrop (Thermo Scientific) and diluted to a concentration of 5 ng/μl. Prokaryotic V4 hypervariable region of 16S rRNA gene (rDNA) was amplified with Phusion High-Fidelity DNA polymerase (ThermoFisher) using universal prokaryotic primers 515F – 806R with further modifications (Corporaso et al., 2012; Parada et al., 2016) using recommended thermocycler conditions with 35 cycles. The universal eukaryotic barcode V9 region of 18S rRNA gene (rDNA) was amplified using 1389F and 1510R primers (Amaral-Zettler et al., 2009), under recommended thermocycler conditions with reduced number of cycles (25) (de Vargas et al., 2015). All amplifications were done in triplicate in order to balance the variance within samples while obtaining adequate amounts of amplified DNA, combined, and then purified using a PCR clean-up kit (Syngen). The final concentration and quality of amplicons were again assessed by NanoDrop, and the library preparation and sequencing experiment on the Illumina MiSeq platform was performed by an external company (Genomed SA). The sequencing yielded 300 paired-end reads targeted for 100 000 reads per amplicon. The raw sequencing data were submitted to the European Nucleotide Archive (ENA) under the accession number PRJEB71447 (ERP156246).

### 2.4 Sequence Analysis

Sequence quality checks were conducted on raw sequence data using FastQC (Andrews 2010), then sequencing adapters and primers were trimmed by trimmomatic (Bolger et al., 2014). Subsequently, processed reads were imported into the qiime2 environment and sequencing primers were removed using cutadapt (Martin, 2011; Bolyen et al., 2019). Finally, DADA2 denoising was done for each sequencing event independently, after which ASV (Amplicon Sequencing Variant) tables and feature tables were merged (Callahan et al., 2016). Taxonomic assignment of V4 16S rDNA was done using an RDP classifier encapsulated in qiime2 against SILVA99 138 database (Wang et al., 2007; Quast et al., 2012), and ASVs classified as ‘Eukaryota’, ‘chloroplast’ or ‘mitochondria’ were discarded. The assignment of V9 18S rDNA ASVs was done by usearch global alignment implemented in vsearch (Rognes et al., 2016) (minimum identity 60% and minimum query coverage 90%) against Protist Ribosomal Database PR2 4.14 (Guillou et al., 2012), prepared following Tara Oceans guidelines (de Vargas et al., 2015). ASVs with the closest hit to a eukaryote, but with an identity lower than 80%, were assigned as an ‘unknown eukaryote’, and the rest were assigned to the best hit. In addition, prokaryotic V9 sequences were classified with usearch global alignment against the SILVA99 138 database (Quast et al., 2012). Before the main analysis, ASVs annotated as Metazoa, Embryophyta, Bacteria or Archaea were also filtered out. Furthermore, we assigned protistan ASVs to one of three trophic groups (‘phototrophic’, ‘consumer’ and ‘parasitic’) based on their taxonomic assignment and the published guidelines (Singer et al., 2021). Additionally, selected ASVs were manually annotated by literature research and assigned to the group ‘mixotrophic’.

### 2.5 Statistical analyses and data visualisation

Statistical analyses, such as alpha-diversity, beta-diversity, metaNMSD, ADONIS, ANOVA or envifit and ancillary data visualisations, were performed in the R environment (version 4.0.3) within RStudio IDE (Allaire, 2012) using packages: vegan (Oksanen et al., 2023), qiime2R (Bisanz, 2018), ggplot2 (Wickham, 2016), phyloseq (McMurdie & Holmes, 2013), ape (Paradis & Schliep, 2019), and microbiome (Lahti & Shetty, 2017). To remove the effect of inequality of sequencing depths, datasets were normalised using scaling with ranked subsampling – SRS (Beule & Karlovsky, 2020). Eukaryotic and prokaryotic prevalence analyses were performed using the microbiome package (Lahti & Shetty, 2017). The analysis was run for sampling sites A1, A2, B1, C1 and C2 (82 samples in total; sample B2 was analysed separately) from the whole sampling season. To compare these datasets, equal ranges of abundance and prevalence thresholds were set and visualised using ggplot2. To investigate synchrony between 18S V9 and 16S V4 rDNA datasets, we used pca-based co-inertia analysis implemented in the ade4 package (Chessel et al., 2004). Two separate analyses were performed: i) for samples representing epilimnion (sites A1, A2, B1, C1, C2), and ii) for the sample from hypolimnion (site B2). Each data table was first SRS normalised to get an even depth for each sample and Hellinger transformed according to (Obertegger et al., 2019). The statistical significance of those analyses was checked with the Monte-Carlo method implemented in RV.test from ade4 with 99 permutations. The PCA-based co-inertia analysis was visualised using ggplot2. To distinguish dead cells coming from upper layers of the lake from potentially living and thriving protistan lineages in anoxic conditions, we compared relative abundances of ASVs in epilimnion and hypolimnion layers during the stratification period (June - August), and only ASVs that achieved more than 2% of relative abundance in at least one time point over this period.

### 2.6 Feature selection by Random Forests and ANCOM-BC

To identify 16S rDNA and 18S rDNA ASVs whose abundance corresponded with seasons (two samplings in the spring, n = 10; three samplings in the summer, n = 15; and three samplings in the autumn, n = 15 per sequencing marker) in samples A1, A2, B1, C1 and C2 (epilimnion and metalimnion), we used supervised machine learning algorithm – Random Forests (Breiman, 2001). This type of method has been proven to perform well in classification of amplicon data (Hermans et al., 2020; Fang et al., 2022). To account for the substantial environmental variability in deeper layers associated with lake mixing, we excluded samples from the deepest point (B2), then ASVs that have more than 0.1% contribution were kept. ASV tables were then re-normalized after the filtration step and transformed using the scale function into scoring units. The data was used to train random forest models and then, Out-Of-Bag (OOB) error was estimated. For each dataset, we picked 30 ASVs with the highest mean decrease Gini coefficient index scores, which corresponded to the highest impact on the classification of the samples, and visualised them with heatmaps and ordination plots. To confirm results from RF, we employed the Analysis of Compositions of Microbiomes with Bias Correction (ANCOM-BC) (H. Lin & Peddada, 2020). The ANCOM-BC was run on absolute counts of the same samples as RF analysis with the option “conserve”, as recommended for the low number of samples, and *p*-values were adjusted using the Bonferroni correction. Subsequently, ASVs which were significantly different (p-value < 0.05) in abundance were compared to ASVs selected by RF. For the visualisation, we also added the 30 most abundant ASVs (estimated based on the sum of reads) for each of the analysed datasets.

## 3. Results

### 3.1 Protist and bacteria diversity in Lake Roś

To investigate the plankton diversity in Lake Roś, we employed V9 18S rDNA and V4 16S rDNA amplicon sequencing of microbial community samples. The samples were collected eight times over a seven-month period, ranging from April 13th to November 18th to represent all the changes occurring during the vegetation season. We collected two size fractions of microorganisms: the prokaryotic fraction (0.2-3 μm) and protists fraction (3-12 μm), from three different sampling locations and two depths, resulting in 96 samples (48 samples per molecular marker). For V9 18S rDNA, we generated a total of 7 628 535 reads (between ∼95 000 and 300 000 per sample) and inferred 7 296 ASVs. Similarly, for V4 16S rDNA, we obtained 5 456 941 reads (between ∼40 000 and 170 000 per sample) and inferred in total of 6 096 ASVs (Supplementary Table S3). The rarefaction curve visualisations for both datasets confirmed that all samples reached the plateau phase (Supplementary Figure S1). For eukaryotic amplicon analysis, we filtered out ASVs assigned to prokaryotes and Metazoa, resulting in a final dataset comprising 5 191 ASVs accounting for 71% of the initial ASV count, with each sample containing approximately 200 to 900 ASVs. It is noteworthy that up to 1% of the prokaryotic sequences filtered out from the V9 18S rDNA datasets were classified as Archaea, including lineages such as Methanosarcina, Nanosalinia, and Nanoarchaeia (Supplementary Figure S2A). Two supergroups (*sensu* Burki et al., 2019) were highly prevalent across all samples: Cryptista (mainly cryptophytes – 24.6% and katablepharids – 3.4%) and TSAR (telonemids – 0.6%, stramenopiles – 21.4%, alveolates – 24.5%, and rhizarians – 6.7%) (Figure 1C). Regarding the V4 16S rDNA dataset, we excluded eukaryotic sequences (predominantly chloroplastic and mitochondrial) resulting in a dataset comprising 5 690 ASVs, representing 93% of the initial count. Each sample contained around 300 to 800 ASVs, and over 99% of these were classified as Bacteria (Supplementary Figure S2B). At the phylum level, the dominant bacterial phyla were Proteobacteria (40.8%), Bacteroidetes (25.2%), and Actinobacteria (24.5%) (Figure 1D). We also observed the temporal changes in the taxonomic composition of the protist (Supplementary Figure S3A) and bacterial (Supplementary Figure S3B) communities at different sites and depths.

### 3.2. Protist and bacterial community structure is shaped by seasonal changes

We observed significant fluctuations in environmental parameters, including water temperature, oxygen levels, light penetration, and the availability of chemical compounds across our sampling events. The surface water temperature varied between 6 and 9°C in April and November, while in June and August, it peaked at 23°C. Notably, the recorded vertical profile of temperature and oxygen concentration during the period from June to August clearly indicated the occurrence of stratification (Supplementary Figure S4). During the summer stratification phase, the deepest sampling location (B2 - 25 m) was below the pronounced oxycline and thermocline, and was characterised by the low temperature (∼10 °C), oxygen deficits and high concentrations of both organic and inorganic compounds in deepest layers (Supplementary Table S2).

Observed changes of the environmental parameters have discernible effects on the composition of microbial communities. In order to assess the diversity and similarity of these communities across various sampling sites and time points, we conducted an analysis of α (richness) and β diversity. Shannon metrics varied between ∼2.3 to ∼5.4 for eukaryotes and between ∼3.5 and 5 for prokaryotes. Notably, if we consider the sampling points, the highest eukaryotic diversity was observed in July at site B, while the lowest occurred in May at the same site (Figure 1E). For the prokaryotic community, peak of diversity was noted in August at site C, with the lowest diversity observed in September at site B (Figure 1F). Alpha diversity for eukaryotic plankton exhibited temporal dynamics, with lower diversity during spring and late autumn, ranging between approximately 3 and 4, and higher diversity during summer, ranging between 4 and 5. This trend was supported by Anova analysis, revealing statistically significant differences between the “month” and “season” categories (p-value < 0.001). Moreover, the differences in eukaryotic α diversity were significant in the geographical microscale and between depths within each month (Supplementary Data 1). The pattern of α diversity for prokaryotes was quite different, however it was also fluctuating – after high diversity in April (∼ 4.5), it dropped during May, June and July (varied between 4 and 4.2) and increased again in late summer and autumn (∼ 4.7). Anova analysis indicated statistically significant differences between months, seasons, sampling sites, and depths within each month (p-value < 0.05) for prokaryotic fraction (Supplementary Data 2). To further understand the diversity of analysed prokaryotic and protist communities, we investigated their β diversity using the Unweighted UniFrac distance metric in conjunction with Multidimensional Scaling (MDS). In both the prokaryotic and protist fractions, we observed that sampling points formed three distinct clusters corresponding to the seasons (spring, summer and autumn) points (Figures 1G and 1H), which was further confirmed with adonis analysis (p-value < 0.001) and beta-dispersion analysis (p-value > 0.05). Noteworthy, in our analysis of β diversity, we did not detect any statistically significant differences between sampling sites for either 18S rDNA or 16S rDNA datasets (Supplementary Data 1 and 2). Due to the formation of hypolimnion at site B2 from June to August (Supplementary Figure S4B), the samples from this spot were considered separately for further analysis.

Despite differences in diversity metrics and the sizes of core communities, the Principal Component Analysis-based Canonical Integration Analysis (PCA-based CIA) indicated that prokaryotic and eukaryotic datasets displayed a coherent pattern of changes across seasons. Analyses have been performed separately for the samples A1, A2, B1, C1 and C2 representing epi-and metalimnion, and the sample B2 representing hypolimnion. The strongest synchrony (RV = 0.96, p-value <0.05) was observed for B2 samples (25 m) compared to samples from the shallower layers (RV = 0.87, p-value <0.05). The synchrony in the upper layers was disturbed in several cases, especially during the summer, when it had a strong variation between prokaryotic and eukaryotic datasets compared to other seasons, which could cause the decrease of RV score (Supplementary Data 3, Supplementary Figure S5).

### 3.3 Dominant and prevalent taxa composition does not correspond with seasonal changes

Observed diversity within both fractions - prokaryotes and eukaryotes accounted for more than 5000 of ASVs. Around 40% of prokaryotic and eukaryotic ASVs were present in every sampling site, and 41.8% of eukaryotic ones and 34.4% of prokaryotic ones were endemic for one spot. The most endemic eukaryotic ASVs were noted in site A (30.7%) with only 14.5% endemic prokaryotic ASVs. On other hand, the highest percentage of prokaryotic endemic ASVs was noted in the site C (15.6%) with only 3% of unique eukaryotic ASVs (Supplementary Figures S6A and S6B).

Therefore, in further analyses of seasonal changes in epilimnion and metalimnion, we focused on the 30 most dominant ASVs for eukaryotic and prokaryotic fractions, which presumably are the main players in the investigated community (Figure 2).

**Figure 2.**
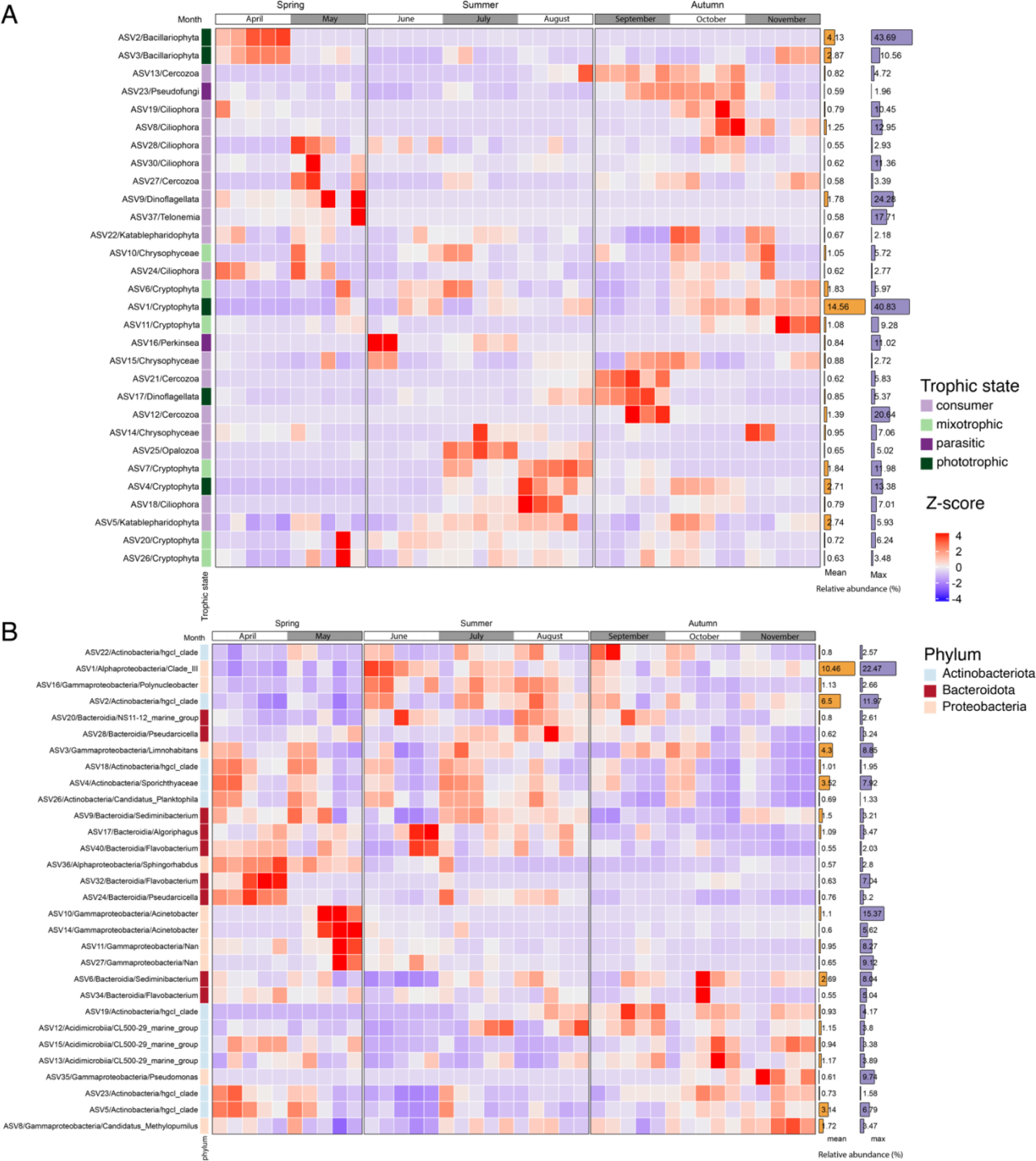
Dominant eukaryotic and prokaryotic ASVs in the epilimnion during the entire sampling period. (A) Heatmap depicting the abundance of 30 dominant ASVs for eukaryotes and (B) heatmap depicting the abundance of 30 dominant ASVs for prokaryotes. The trophic states for eukaryotes and phyla for prokaryotes are colour coded.

Among the dominants (30 ASVs), the most persistent eukaryotic ASVs mainly belonged to the mixotrophic cryptophytes (ASV1, ASV6, ASV20, ASV26) and chrysophytes (ASV10), as well as to the predatory katablepharidophytes (ASV5, ASV22), ciliates (ASV24, ASV28) and cercozoans (ASV27). In addition, we were able to identify separate assemblages of ASVs associated with the particular seasons throughout the sampling period. In spring, we observed a high abundance of diatoms (ASV2 and ASV3) in April, while ciliates (ASV28, ASV30), predatory dinoflagellates (ASV9) and telonemids (ASV37) were abundant in May. In summer, especially in July and August, we observed the cluster of ASVs, which included opalozoans (ASV25), two cryptophytes (ASV4 and ASV7) and ciliate (ASV18). Autumn was characterised by the increase in abundance of consumers and mixotrophs such as chrysophytes (ASV15), cercozoans (ASV12, ASV21) and dinoflagellates (ASV17) in September and the increase in abundance of ciliates (ASV8, ASV19), cercozoans (ASV13) and a member of the parasitic pseudofungi (ASV23) in October and early November (Figure 2A).

Similar to eukaryotic plankton, most bacterial ASVs persisted throughout the sampling period, but we could identify differences in their abundance between seasons. The dominant ASVs belonged to three phyla – Actinobacteriota (11), Bacteroidota (9) and Proteobacteria (10). Seven ASVs were more abundant in spring compared to the other seasons. Three of them dominated in April – the genus *Sphingorhabdus* (Alphaproteobacteria; ASV36) and the genera *Flavobacterium* (ASV32) and *Pseudarciella* (ASV24), which belong to the Bacteroidia. In May, two *Acinetobacter* ASVs (ASV10, ASV14) as well as ASV11 and ASV27, which all belong to the Gammaproteobacteria, dominated. Most ASVs were associated with either the summer or autumn months. In summer, members of hgcI clade (Actinobacteria; ASV2), clade III (Alphaproteobacteria; ASV1) and *Polynucleobacter* (Alphaproteobacteria; ASV16) as well as ASVs of different Bacterioidia genera (ASV17, ASV20, ASV28, ASV40) reached the highest relative abundances. In autumn, we observed an increased abundance of ASVs assigned to three members of the CL500-29 marine group (Acidimicrobiia; ASV12, ASV13, ASV15), two members of the hgcI clade (ASV19, ASV22), two Gammaproteobacteria– Candidatus Methylophilus (ASV8) and *Pseudomonas* (ASV35) - and two taxa assigned to Bacteriodia (ASV6, SV34). The abundance of seven ASVs remained fairly constant across all seasons, including five ASVs belonging to the hgcI clade (ASV4, ASV5, ASV18, ASV23, ASV26) and two ASVs representing the genera *Limnohabitans* (Gammaproteobacteria; ASV3) and *Sediminibacterium* (Bacteriodia; ASV9).

Even though microbial communities change dramatically between seasons, we expected some ASVs to be present in high numbers during the whole season. To identify the cosmopolitan protists and bacteria in the surface layer (epilimnion) we applied the prevalence analysis. Only several eukaryotic ASVs (16) were highly prevalent (occurred in more than 70% samples) and therefore could be considered as ‘core’ microbiome for the ice-free season (Supplementary Figure S6C). Those belonged to Hacrobia (7), Alveolata (5) and Stramenopila (4), and accounted, on average, for 31% of relative abundance of all protists. Only two ASVs were present in every analysed sample - ASV1 (classified to the genus *Cryptomonas*) and ASV5 (Katablepharidophyta) (Supplementary Figure S6E). In contrast, the core prokaryotic microbiome was much more numerous (50 ASVs) and accounted, on average, for 54% of the relative abundance of prokaryotes (Supplementary Figure S6D and S6F). Moreover, the ratio between the mean and the maximum relative abundance of protist (“division” level) and prokaryotic (“class” level) taxa were much higher for eukaryotes (maximum 19-fold, noted for Discoba) than prokaryotes (maximum 7-fold) (Supplementary Tables S4 and S5). Such differences suggest higher variability of the abundance of protists than prokaryotes over the vegetation season.

### 3.4 Random Forest analysis unveiled pivotal ASVs for seasonal community structures

The dominant ASVs analysis did not explain well observed dynamic changes in microbial community structures across different seasons. To address this, we employed supervised machine learning (Random Forests) and statistical (ANCOM-BC) methods, to identify ASVs significantly contributing to the shifts in community structure between seasons. For each dataset, we focused on the top 30 ASVs deemed most significant by the Random Forest (RF-selected) model, as determined by the mean decrease Gini coefficient indexes (Supplementary Figure S7). Through the implementation of RF models, we effectively categorized our samples into three distinct seasons (with an out-of-bag (OOB) estimate error rate of 0% for the 18S rDNA dataset and 2.5% for the 16S rDNA dataset). The ASVs revealed selected by RF were also identified as statistically significant by the ANCOM-BC models, however only 5 ASVs for eukaryotes and 2 ASVs for prokaryotes exhibited overlap with the 30 most abundant ASVs (Supplementary Figure S8).

The 30 eukaryotic ASVs identified by RF analysis showed a non-uniform distribution of taxa across the sampling period, with ASVs clustered not only for the three seasons but also for the months (Figure 3A). Each of these clusters included ASVs representing different trophic states, such as phototrophs or mixotrophs, consumers and parasites, which were assessed based on literature searches (Supplementary Table S6). Spring was represented by twelve rf-selected ASVs. The presence of Bacillariophyta (ASV157) and parasitic fungi (ASV32) – chytrids - was consistent with the diatom bloom that typically occurs in April. During this period, we also observed a high abundance of photosynthetic Eustigmatophycae (ASV46; *Nannochloropsis*) and ASV59, which are assigned to predatory *Stoeckeria* (Dinoflagellata). In May, most ASVs belonged to the heterotrophic assemblage, which consisted of ASVs representing dinoflagellates (ASV9), ciliates (ASV31, ASV85), cercozoans (ASV63, ASV68 and ASV113) and chrysophytes (ASV114), while parasites were represented by the genus *Lagenidium* (ASV164; Pseudofungi, Stramenopiles). In summer, the RF model and ANCOM-BC indicated the turnover of the observed taxa compared to the composition in spring. We were able to identify a more diversified group of photosynthetic or mixotrophic taxa belonging to chlorophytes (2), woloszynskioid dinoflagellates (2), Raphidophyceae (1) and Cryptophyta (1). Heterotrophs were also diverse, including Ciliophora (3), Stramenopila – MAST-12 (1), Centroheliozoa (1) and Choanoflagellata (1). An ASV representing parasites was also observed (ASV16; Perkinsea). Moreover, taxa associated with summer were not evenly distributed, with a clear shift in the main ASVs assigned as primary producers and consumers between months. Among phototrophs, chlorophytes (ASV45; ASV100) reached high relative abundance in June, woloszynskioid dinoflagellates (ASV133; ASV187) together with ASV111 (Raphidophyceae) in July, and ASV4 (Cryptophyta) in August. A similar succession was observed in the consumers: ciliate (ASV49), choanoflagellate (ASV80), MAST-12 (ASV65) and Chrompodellida (ASV149) were particularly conspicuous in June, a centroheliozoan (ASV130) and a ciliate almost identical to *Halteria* (ASV124) in July, while only a single ASVs (ASV85; Ciliophora) was noticeable in August. Autumn can be roughly divided into two periods, the first (September and some spots in October) being characterised by a high relative abundance of various Cercozoa (ASV12, ASV13, ASV68 and ASV113). In the second period (October and November), various consumers were present, which were assigned to Cercozoa (ASV63), Stramenopiles (*Labyrinthulea*; ASV132) and Amoebozoa (ASV126). Phototrophs, which were particularly abundant in October, were assigned to the Bacillariophyta (ASV33), Cryptophyta (ASV4) and Eustigmatophyta (ASV46). However, some of the ASVs that were clearly associated with a particular season were also present in other seasons. For example, the aforementioned ASV46, which was assigned to the Eustigmatophyta, was mainly present in early spring and late autumn, or the Cercozoa (ASV113), whose relative abundance was high in either May or September. Grouping by season and individual months was also supported by the Bray-Curtis MDS visualisation, which contrasts with the Bray-Curtis MDS visualisation of dominant ASVs, where data points were clustered together except for those from April (Supplementary Figure S9A and S9C).

**Figure 3.**
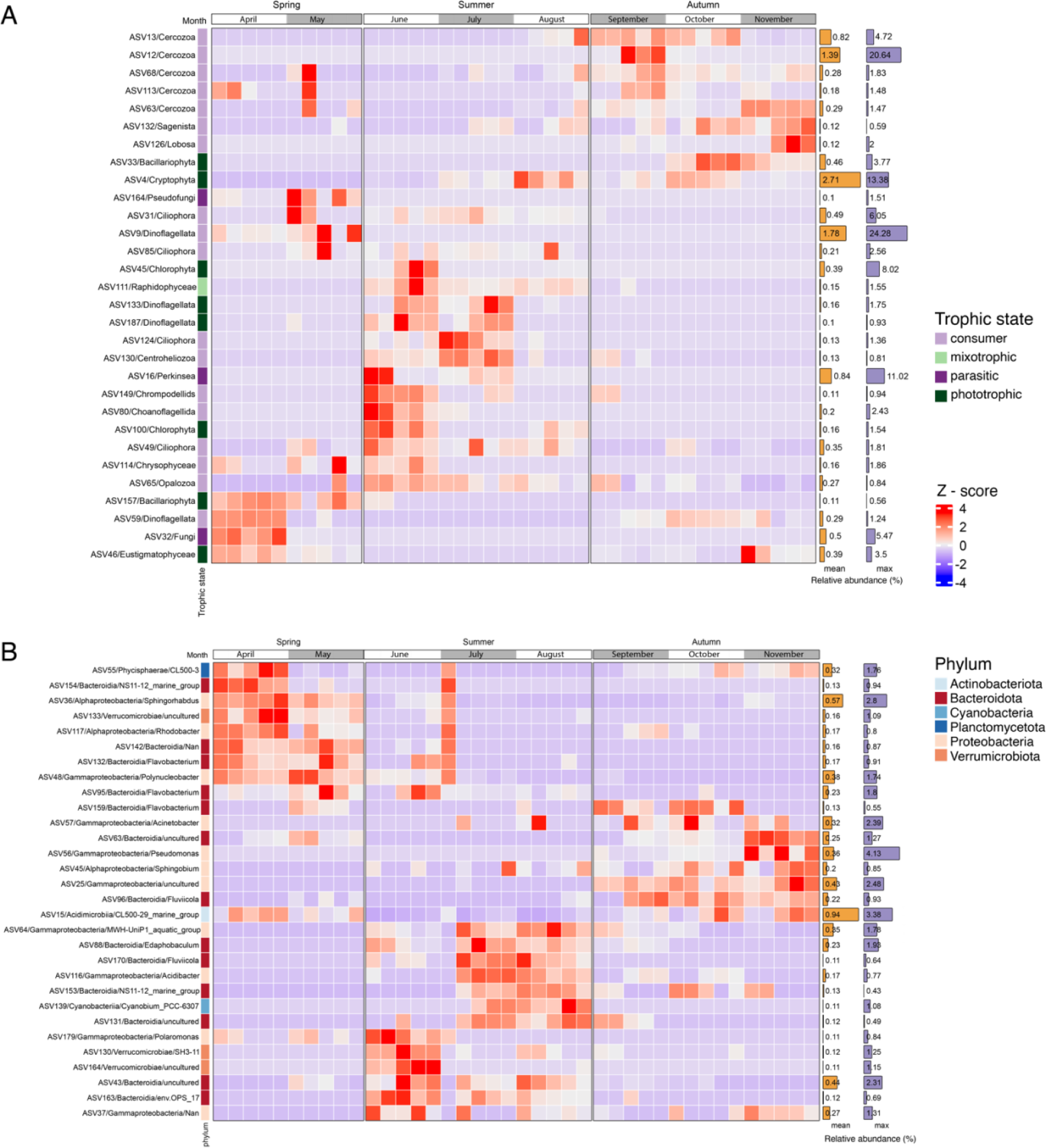
Model selected eukaryotic and prokaryotic ASVs in the epilimnion throughout the entire sampling period. (A) Heatmap depicting the abundance of 30 ASVs selected by the random forest model for eukaryotes (B) Heatmap depicting the abundance of 30 ASVs selected by the random forest model for prokaryotes. The trophic states for eukaryotes and phyla for prokaryotes are colour coded.

Analysis of the RF-selected ASVs within the prokaryotic dataset also revealed a non-uniform distribution of taxa with four main groups of ASVs assigned to six phyla – Actinobacteriota (1), Bacteroidota (13), Cyanobacteria (1), Planctomycetota (1), Proteobacteria (11) and Verrumicrobiota (3) (Figure 3B). The first group of ASVs was associated with spring, with a considerable presence of ASVs assigned to the Bacteroidia (ASV132, ASV95, ASV142, ASV154), two genera of Alphaproteobacteria – *Sphingorhabdus* (ASV36) and *Rhodobacter* (ASV117), ASV48 (*Polynucleobacter*, Gammaproteobacteria), Verrucomicrobiae (ASV133) and CL500-3 clade of Phycisphaerae (ASV55). Moreover, five of them occurred mainly in April (ASV55, ASV154, ASV36, ASV133 and ASV117), while four of them reached higher abundance in May (ASV142, ASV132, ASV48, ASV95). In summer, we observed two separate clusters of ASVs – the first in June and the second in July and August. The ‘early summer’ grouping consisted of six ASVs – classified as Bacteroidia, assigned to the env.OPS 17 group (ASV43 and ASV163), Verrucomicrobiae (ASV130 and ASV164) and Gammaproteobacteria (ASV37, ASV179). The ‘late summer’ cluster of ASVs was formed by seven ASVs belonging to Bacteroidia (ASV88, ASV131, ASV153, ASV170), *Cyanobium* PC-6307 (Cyanobacteria, ASV139) and Gammaproteobacteria (ASV64, ASV116). Interestingly, in a single sample in July (A1), we observed the recovery of the spring-associated assemblage of ASVs. In autumn, ASVs were inconsistently distributed, with some ASVs being more abundant at the beginning (September and October) such as ASV159, assigned to Flavobacterium (Bacteroidia), and ASV57 (*Acinetobacter*, Gammaproteobacteria) or at the end (October and November) such as members of Gammaproteobacteria (ASV56 and ASV57) and Bacteroidia (ASV63). However, ASV45 (*Sphingobium*, Alphaproteobacteria) and ASV96 (*Fluviicola*, Bacteroidia) persisted throughout the whole autumn. In addition, there were ASVs that could not be explicitly associated with a specific time of the sampling period, such as ASV15 (Acidimicrobiia), which reached higher relative abundances in both spring and autumn, or ASV37, which was present in numerous samples in summer and autumn. The Bray-Curtis MDS visualisation of the samples consisting of RF prokaryotic ASVs showed a slightly different arrangement to that of the eukaryotic ASVs, with three distinct clusters representing the seasons, albeit without a smooth transition between months. This is also in contrast to the Bray-Curtis MDS visualisation of the prokaryotic dominant ASVs, where all data points were clustered together (Supplementary Figure S9B and S9D).

### 3.5 Distinct microbial community is established in the deep lake waters throughout the summer months

Throughout the summer months, spanning from June to August, Lake Roś undergoes a period of stratification, marked by the presence of a distinct thermocline that separates the epilimnion and hypolimnion layers, each exhibiting markedly different environmental conditions. In comparison to the surface water, the hypolimnion layer was characterised with low temperature (∼ 10°C) and the absence of sunlight and oxygen. Beta-diversity analyses of prokaryotic and eukaryotic plankton revealed that during this period, a discrete cluster of samples emerged within the hypolimnion layer at site B, located at a depth of 25 meters (Figure 4A) implying the presence of a distinct microbial community. In contrast, dissimilar microbial communities at different depths were not observed at other sampling sites, denoted as A and C (with deeper sampled fractions at 6 and 10 metres), despite these locations also exhibiting episodes of anoxic conditions during this period (Supplementary Table S2), suggesting that the distinct community of the site B is not only shaped by lack of oxygen.

**Figure 4.**
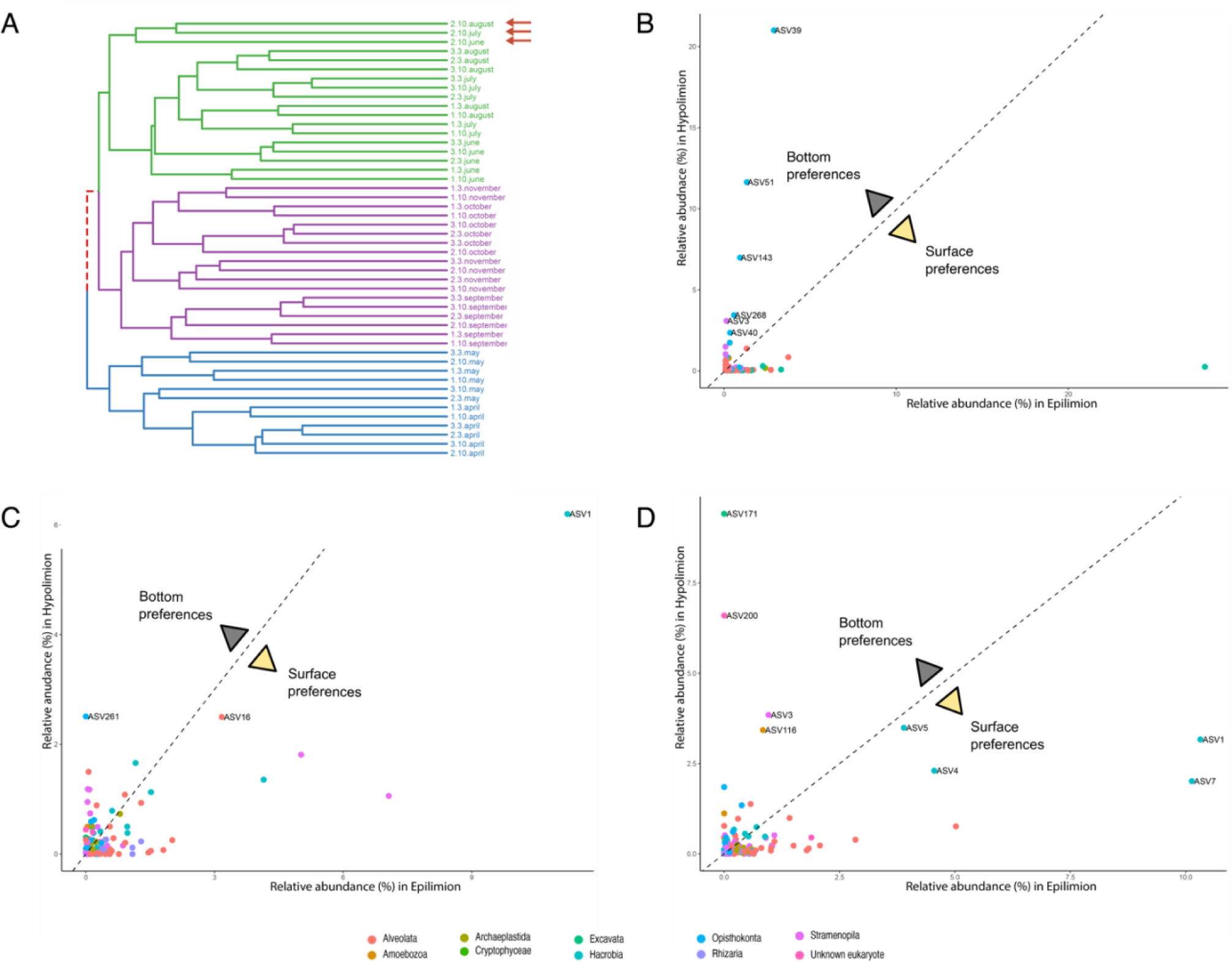
Preferences of eukaryotic taxa between the hypolimnion and the epilimnion during stratification at site B2 (25 m). (A) Dendrogram of the eukaryotic samples (18S V9 rDNA) based on the Unweighted UniFrac metric (“complete” clustering method), with the hypolimnion samples marked with red arrows. Comparisons of relative abundances (%) of ASVs between the surface (B1) and bottom part (B2) for June (A), July (B) and August (C). ASVs that reached more than 2% of relative abundance in a given sample were labelled. Eukaryotic groups are colour coded.

An analysis of the taxonomic comparison of eukaryotic ASVs between the epilimnion and hypolimnion during the stratification period confirmed the differences in community structure (Supplementary Figure S3). However, rather than observing a consistent community structure persisting throughout this period, we unexpectedly observed the formation of distinct assemblages for each of the three individual months (Figure 4). In June, five fungal ASVs achieved highest relative abundances (45%), accompanied by a single diatom ASV assigned to *Stephanodiscus* (ASV3) – 3% (Figure 4B; Supplementary Table S8). At the same time, only one ASV classified as Cryptophyta (ASV1) dominated the epilimnion (28%). In July, only two ASVs were noted with high relative abundances – the previously described diatom ASV3 (6%) and ASV261 (2.5%) annotated as choanoflagellate (Figure 4C; Supplementary Table S8). Finally, the hypolimnion layer in August was mainly inhabited by a bodonid (ASV171) – 9.5%, and *Vermamoeba* (ASV116) – 3.5%, followed by an unknown eukaryote (ASV200) – 7%, diatom (ASV3) – 4%, and a representative of Katablepharidophyta (ASV5) – 3.5% (Figure 4D; Supplementary Table S8). An analysis of the distribution of ASVs associated with the hypolimnion revealed several protistan ASVs (Supplementary Table S8) across all samples, strongly implying their association with anoxic water conditions. The exception was ASV116 (identified as *Vermamoeba*), which was exclusively found in site B2 at a depth of 25 meters (Supplementary Figure S10A).

The prokaryotic community structure was more uniform during the stratification period with a prevailing presence of two phyla – Proteobacteria and Bacteroidota (Figure 5A, B, C). An analysis of the distribution of ASVs associated with the hypolimnion, at the taxonomic level of “Family” across all samples, revealed a notably higher abundance of these ASVs in sites B2 and C2 in comparison to the other sampling locations (Supplementary Figure S10B).

**Figure 5.**
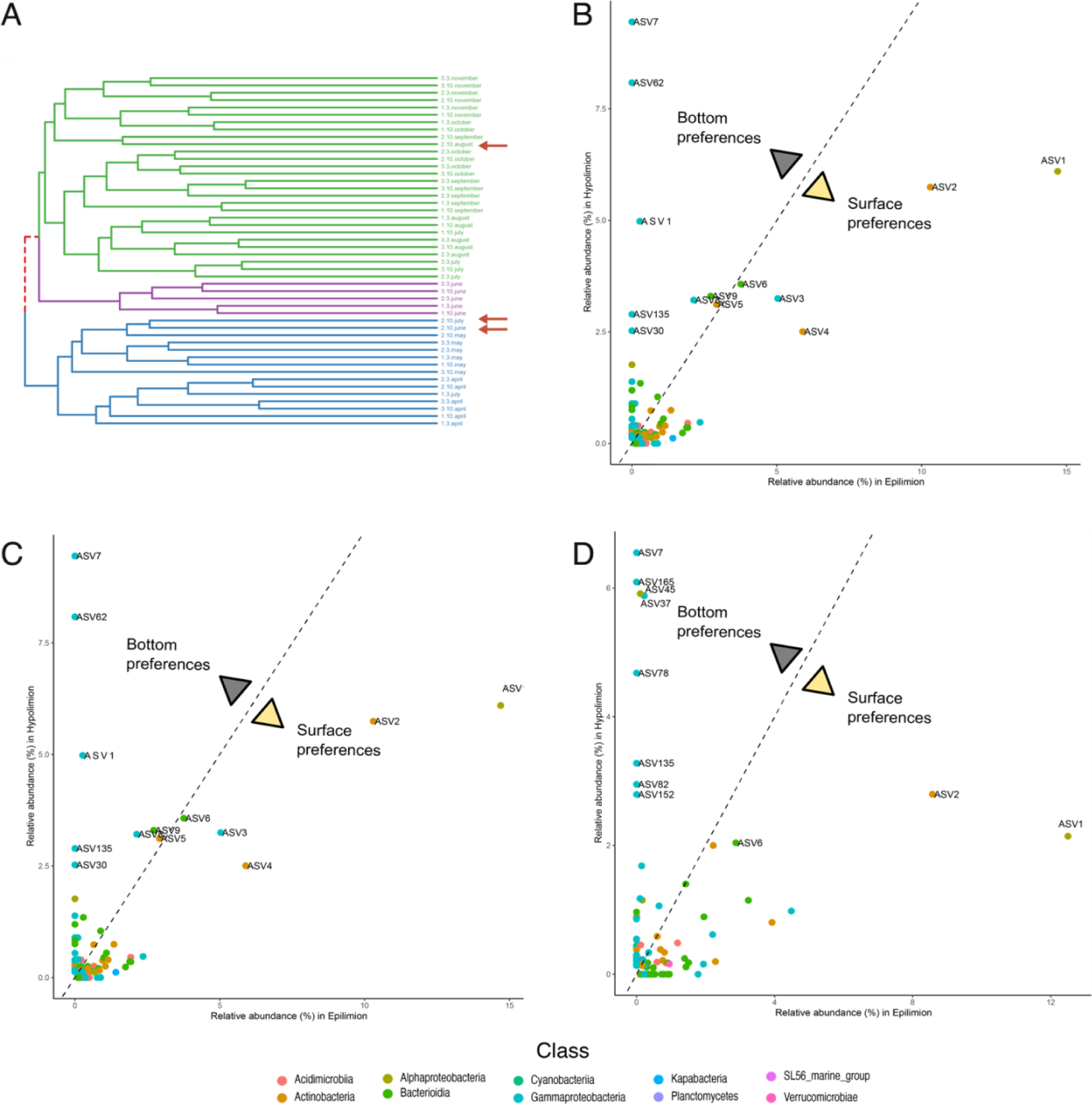
Preferences of prokaryotic taxa between the hypolimnion and the epilimnion during stratification at site B2 (25 m). (A) Dendrogram of the prokaryotic samples (16S V4 rDNA) based on the Unweighted UniFrac metric (“complete” clustering method), with the hypolimnion samples marked with red arrows. Comparisons of relative abundances (%) of ASVs between the surface (B1) and bottom part (B2) for June (A), July (B) and August (C). ASVs that reached more than 2% of relative abundance in a given sample were labelled. Prokaryotic classes are colour coded.

### 3.6 Abiotic and biotic factors influenced the microbial community composition

Through the incorporation of environmental parameters into our analyses, we were able to discern the factors that exerted influence on the observed gradient or continuum of communities, as indicated by their correlation with the Non-Metric Multidimensional Scaling (NMDS) axes (Figure 6A, B). In total, 18 environmental factors were tested using the envfit function, which entails fitting environmental vectors onto the NMDS ordination plot. This analysis revealed that nine factors were significantly correlated with the ordination (p-value <0.01) for eukaryotic communities, and 15 factors exhibited significant correlations with bacterial communities. The water and the air temperature, oxygen and Si concentration were the main factors shaping the structures of both communities. Other parameters such as light penetration, Secchi disk visibility, concentrations of NO_2_, NO_3_, NH_3_, Mn, TC, TN, Fe, P, and the Trophic State Index (TSI) were congruent with the prokaryotic community structure. Dissolved organic matter (DOC) was found to be a significant driver exclusively for the eukaryotic community structure.

**Figure 6.**
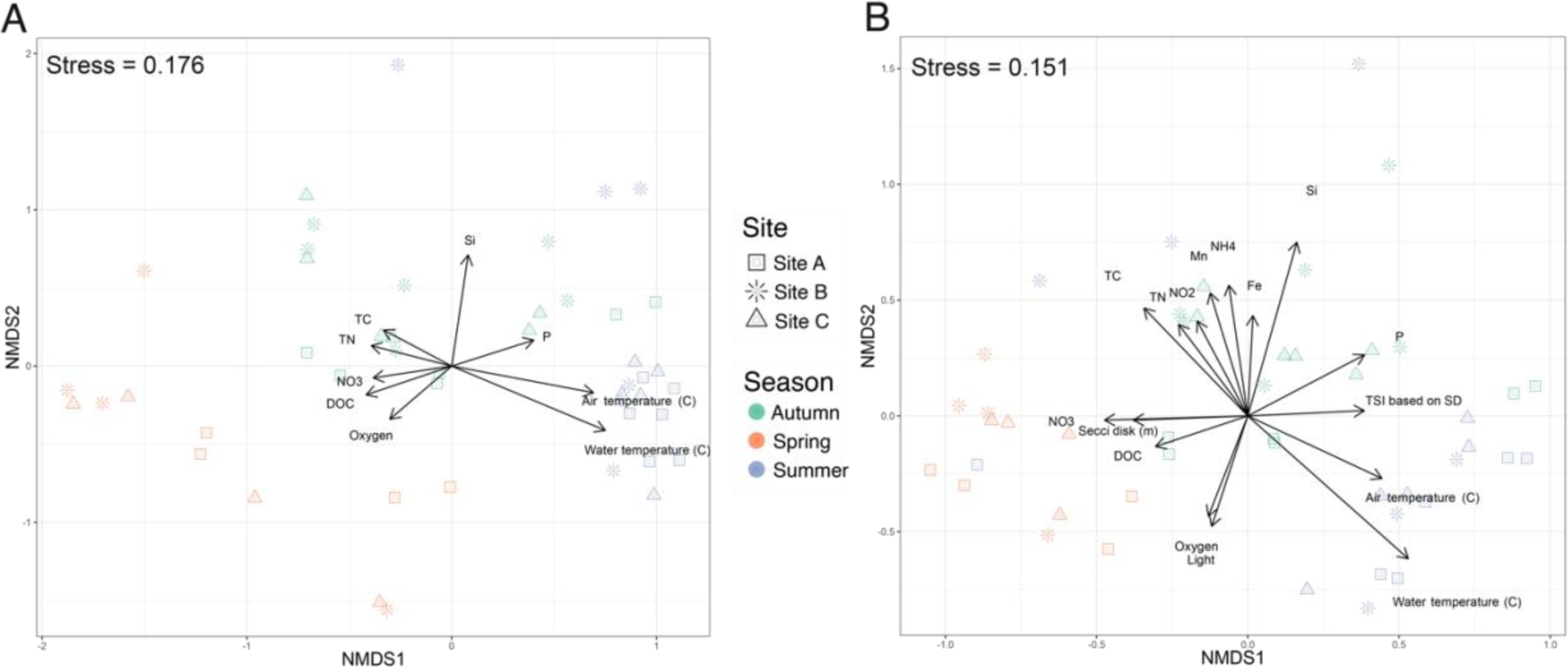
Impact of physicochemical factors on microbial community structure. Sample composition of (A) eukaryotic and (B) prokaryotic datasets based on NMDS analysis with envfit (vegan package) fitted physicochemical parameters (p <0.05). The colours refer to the seasons, the shapes to the sampling sites.

The influence of abiotic factors on microbial community structure is profound; however, at specific time points, biotic factors appear to assume a critical role. Our focus centered on the grazing impact of zooplankton on the protist community, and we conducted an analysis of the presence and abundance of the principal zooplankton groups. We used microscopic data collected for eleven zooplankton groups to perform abundance comparisons and correlation analyses and reveal their interactions with prokaryotic and eukaryotic plankton communities. The abundance of zooplankton displayed temporal dynamics throughout the year, with the highest numbers occurring during the spring (May) and autumn (October) (Supplementary Figure S11). The marked increase in zooplankton abundance and grazing activity can explain an unexpected clustering pattern of protist groups across sampling sites within the same time points (Supplementary Figure S12). Such unexpected pattern was observed in the samples from May, where sample C1 exhibited a close relationship with B2, while sample C2 clustered with sample B1. Within the C2-B1 cluster, a notably high relative abundance of dinoflagellates (23%/31%) and ciliates (12%/30%) was observed, with a limited abundance of cryptophytes. In contrast, the B2-C1 cluster was dominated by cryptophytes (48%/70%), while ciliates (3%/13%) and dinoflagellates (∼2% in both cases) were less prevalent. The influence of zooplankton on planktonic protists was evident not only at the level of single ASVs but also at the community level. The analysis of the NMDS ordination plot with vectors fitting using envfit (p-value < 0.01) implies that the larvae of Copepoda (nauplii), *Bosmina longirostris*, *Chydorus sphaericus, Leptodora kindtii*, and adult Cyclopoida. were the main drivers for shaping the community structure (Supplementary Figure S13). Spearman’s correlation analysis showed a more substantial impact of zooplankton on the protist community compared to the prokaryotic one, with 137 eukaryotic and 37 prokaryotic ASVs involved in statistically significant correlations (p-value < 0.05 and r2 > |0.4|) (Supplementary Figure S14). The analysis pointed to ciliates (33), chlorophytes (13), dinoflagellates (10) and cercozoans (10) as the taxa whose abundance correlates with zooplankton groups identified as *Asplanchna* sp. (44) and *Diaphanosoma brachyurum* (44).

## 4. Discussion

### 4.1 Diversity of eukaryotic and prokaryotic plankton in Lake Roś

The 16S rDNA and 18S rDNA metabarcoding allowed us to identify the vast diversity of planktonic prokaryotic and eukaryotic community of the Lake Roś. The prevalence of Cryptophyta and Katablepharidophyta (Hacrobia), Ciliophora and Dinoflagellata (Alveolata), and Bacillariophyta (Stramenopila) among eukaryotes (Figure 1C), and Actinobacteria, Bacteroidota, and Proteobacteria among prokaryotes (Figure 1D), was in line with previous reports on the microbial community composition in temperate zone lakes (Liu et al., 2015; Kiersztyn et al., 2019; Cruaud et al., 2019; Cruaud et al., 2020).

We have also observed dynamic changes in the taxonomic composition of both prokaryotic and protistan community across the seasons, indicating a temporal succession of species. This well-known phenomenon finds support in both richness and beta-diversity analyses of the Lake Roś community (Figure 1E-H). Even though the bacterial and eukaryotic communities in Lake Roś exhibited dynamic changes over time in different scale, the co-inertia analysis based on PCA revealed their synchrony, which might be a result of direct biotic interactions or response to the same environmental factors (Bock et al., 2018). This finding aligns with previous metabarcoding studies conducted in various aquatic ecosystems, including rivers, reservoirs, and lakes, which have consistently demonstrated synchronous behaviour between bacterial and protist communities (Kent et al., 2007; Bock et al., 2018; Bock et al., 2020; Mikhailov et al., 2022). In Lake Roś, the most pronounced synchrony was observed in the deeper waters (B2), whereas synchrony in shallower waters experienced disruptions on several occasions, particularly during the summer season. These disturbances could be due to factors such as the strong dominance of certain taxa, selective grazing of zooplankton and climatic disturbances such as heavy rainfall and additional mixing in the shallower parts of the lake (Supplementary Figure S5, Supplementary Table S2).

While it is undebatable that the geographical distance has impact on the protist community structure (Schiaffino et al., 2016; Boenigk et al., 2018), much less attention was put into diversity within single water body. Most of the recent studies, whether focused on single or multiple lakes, have characterized microbial communities based on a single sampling site per lake (Simon et al., 2015b; Sieber et al., 2020). That approach, which disregards the local variations in biotic and abiotic factors within lakes and their impacts on microbial community diversity, might lead to some oversimplifications or provide biased assessment. For instance, in the case of Lake Roś, our sampling strategy encompassed three distinct sampling sites, enabling the detection of variations in microbial community diversity within a single lake (Supplementary Figure 5). The statistically significant effect of the sampling site was also observed for much larger fresh water bodies, such as Lake Baikal (David et al., 2021). Although beta-diversity analysis highlighted seasonality as the predominant factor shaping the communities, with the “site” factor not attaining statistical significance, we observed disparities between the core taxa composition in surface layers of sites A, B, and C. Particularly noteworthy were the eukaryotic core communities at site A, representing 30.7% of all ASVs. The prokaryotic community exhibited lower variability, yet the most distinct communities were discerned at sites A and C, constituting 14.5% and 15.6%, respectively. These findings suggest that the microbial community response to local physicochemical and biological factors is significantly influenced by the hydrological characteristics of the lake. For instance, site A was the shallowest sampling site (6 m deep) experiencing consistent year-round mixing, separated from other points by a narrow channel. In contrast, site C is potentially impacted by inflows of allochthonous matter carried by two small rivers that collect water from surrounding farmlands.

The taxonomic composition and seasonal patterns observed in the analysed microbial community of Lake Roś exhibited similarities to findings from other studies conducted in temperate lakes. This congruence provided confidence that Lake Roś could be considered a representative example of a dimictic temperate zone lake. As a result, we were able to delve deeper into the community structure and draw general conclusions on the microbial communities of dimictic lakes. Additionally, our comprehensive sampling strategy, which encompassed three distinct sampling locations within the same lake, allowed us to untangle overarching universal patterns from local variations, including those observed in the lake’s deepest layers. These disparities at the microscale underscore the importance of carefully selecting sampling locations, as it can significantly impact the accurate identification of distinct microbial communities.

### 4.2 Seasonal dynamics of microbial communities in epilimnion and metalimnion

Our primary objective was to examine the seasonal and spatial variations in the microbial community within the context of the dimictic nature of the studied lake, with a particular focus on examining individual ASVs. Microbial communities change over multiple timescales (i.e., from hours, days or weeks to seasons) in response to a multitude of abiotic and biotic factors (Fuhrman et al., 2015). In temperate lakes, the recurring seasonal patterns strongly influence both prokaryotic (Crump & Hobbie, 2005; Rusak et al., 2009) and eukaryotic (Simon et al., 2015b) community composition. However, these changes are often examined at the level of functional groups, encompassing primary producers and consumers, or large taxonomic groups. Our metabarcoding analysis supports previous findings on the seasonal changes, such as PEG model (Sommer et al., 1986; Sommer et al., 2012). However, our study advances further on these findings by using machine learning and statistical methods to provide new and detailed insights into the succession of protist and bacterial ASVs in Lake Roś.

Essentially, we found that the dominant eukaryotic ASVs during most of the ice-free season belonged mainly to the mixotrophic groups (especially cryptophytes and chrysophytes) and eukaryovours katablepharids (Figure 2A, Supplementary Table S8). The continuous presence and predominance of these groups suggests that they do not have a significant influence on the formation of unique seasonal communities. Nevertheless, it is worth noting that the previously mentioned dominant protistan plankton groups could potentially undergo changes during winter, as previous studies with annual sampling have shown (Cruaud et al., 2019). Mixotrophic plankton, particularly phago-mixotrophic organisms that combine photosynthesis with phagotrophy, are of particular interest due to their dual role as both producers and consumers (Millette et al., 2023). On the one hand, their mixotrophic potential can be an advantage and increase their flexibility in adapting to changing environmental conditions by grazing on bacteria and thus displacing phototrophic species (Selosse et al., 2017). On the other hand, increased nutrient and organic matter inputs to certain lakes may affect mixoplankton by promoting bacterial prey population growth and limiting light availability due to humic substance absorption in the water column (Wilken et al., 2018). Laboratory experiments also suggest that mixoplankton may rely more on phagotrophic processes as temperatures rise (Wilken et al., 2013; Lin et al., 2018; Cabrerizo et al., 2019), highlighting the growing importance of mixoplankton metabolism with the warming of lake waters. Other groups of phototrophic and heterotrophic eukaryotic ASVs were not as prevalent and were present in certain periods of the sampling season. Phototrophs were abundant in nutrient-richer colder periods such as April when diatoms dominated, signifying the onset of the spring bloom, or a dinoflagellate (ASV17) in September probably as a result of being outcompeted by mixotrophs during summer nutrient limitation (Figure 2A, Supplementary Table S8). Quite importantly, patchy distribution of predatory protists such as ciliates, cercozoans, telonemids or dinoflagellates clearly pinpoints their different feeding strategies and prey preferences (e.g. based on size), or competition between them (Supplementary Table S7).

In contrast, the bacterioplankton community remained relatively consistent throughout the study period. The group of dominants comprised various representatives of Actinobacteria, Acidimicrobiia, Gammaproteobacteria and Bacteroidota (Figure 2B), previously described as a major component of freshwater ecosystems (Kiersztyn et al., 2019; Cruaud et al., 2020). The only ASVs with a very specific abundance pattern were four ASVs belonging to the Gammaproteobacteria (ASV10, ASV14, ASV11 and ASV27), two of which were assigned to the genus *Acinetobacter*. Although there are not many reports on the seasonality of this group in freshwater systems, their abundance in the majority of samples from May might indicate that they are directly or indirectly involved in ongoing clear water phase, for instance as members of the pathobiome of abundant specimens of zooplankton or fish (Supplementary Figure 11) (Sevellec et al., 2014; Eckert et al., 2016).

### 4.3. Less abundant taxa have a significant impact on seasonal protist communities

To identify the ASVs which contribute the most to the formation of distinct microbial communities in different seasons, we employed RF models and Ancom-BC, which both highlighted the role of less abundant lineages characteristic for each season (Figure 3). In fact, the set of ASVs identified by RF was very different from the dominant ones. The disparity was especially profound for eukaryotes, pointing out possible ecological importance of lineages like phototrophic and heterotrophic dinoflagellates, phototrophic eustigmatophytes and chlorophytes, or heterotrophic cercozoans and ciliates. Importantly, each season was represented by different lineages of protists (different ASVs), even though the same functional categories such as phototrophs, consumers and parasites were present through the whole sampling period (Figure 3A). Among primary producers, beside representatives of expected groups such as diatoms and chlorophytes, we identified Eustigmatophyceae (*Nannochloropsis*), which were previously documented in spring blooms of freshwater lakes (Fawley & Fawley, 2007). Through taxonomic classification and extensive literature searches, we were able to classify some of the detected dinoflagellates as primary producers, particularly those associated with photosynthetic genera *Asulcocephalium* (ASV133) and *Leiocephalium* (ASV187) (Takahashi et al., 2015). The diversity of consumers, on the other hand, was significantly greater in all seasons. We were able to pinpoint taxa essential for grazing within the eukaryotic community. This included two dinoflagellates related to *Gyrodinium* (ASV9) and *Stoeckeria* (ASV59), as well as a ciliate from the *Balanion* genus (ASV49) in the spring. *Gyrodinium* appears to be especially crucial, as it has been previously reported as a major grazer of diatoms in marine systems, as opposed to ciliates, which are less capable of consuming large prey (Saito et al., 2006). Other highly abundant predators in May that may be involved in controlling the decline of diatom blooms were three eukaryovorous cercozoans (Supplementary Table S7) belonging to two closely related genera – *Protapis* (ASV68) and *Cryothecomonas* (ASV13 and ASV63), which are mainly known as typical marine diatom predators (Drebes et al., 1996; Schnepf & Kühn, 2000; Moustaka-Gouni et al., 2016).The remaining key heterotrophs were involved in bacterial grazing. In the spring, ASV114 (*Pedospumella*, Chrysophyceae) likely played a significant role as one of the most important bacterioplankton predators in freshwater ecosystems (Šimek et al., 2013). However, during the summer, bacterivorous protists displayed greater taxonomic diversity, including “rare taxa” representing colpodellids, stramenopiles, heliozoans, and choanoflagellates. This discovery of rare taxa underscores their importance for the summer microbial community and strongly suggests their seasonality (Schiwitza et al., 2020; Zagumyonnyi et al., 2022). Nevertheless, further research is required to investigate their ecological roles in lakes. Among the heterotrophic organisms, we also identified potential decomposers, namely the ASV132, related to *Thraustochytrium* sp. (Labyrinthulomycetes, Stramenopila), which represents a significant group of marine and freshwater saprotrophic eukaryotes known for their ecological role as decomposers (Pan et al., 2017; Morabito et al., 2019; Xie et al., 2022). While we can only speculate on the exact role of this lineage in freshwater ecosystems, it likely contributes to the decomposition of biomass from ongoing summer cyanobacterial blooms. The occurrence of chytrids (ASV32), perkinsids (ASV16), and pseudofungi (ASV164, ASV23), in the seasonal protist communities not only underscores their significant influence on shaping the diversity and dynamics of freshwater ecosystems (Mangot et al., 2009), but also raises concerns for host-parasite interactions which might be impacted with the increase of eutrophication (Budria, 2017). Of particular note is the remarkable diversity of Perkinsea-related ASVs, comprising a total of 106 distinct ASVs. This parasitic group, with a wide host range spanning from dinoflagellates to animals, poses a potential risk in freshwater environments, where it has been linked to the occurrence of mass mortalities among amphibian species (Isidoro-Ayza et al., 2017; Itoïz et al., 2022). The majority of ASVs that play a pivotal role in shaping seasonal communities were exclusively present in a single season. However, there were certain taxa that recurred across multiple seasons, implying a potential adaptation to specific biotic and abiotic factors, such as the presence of prey, nutrient availability, or favourable temperature conditions. For instance, the repeated presence of eukaryovorous dinoflagellates (ASV9; ASV59) could be linked to their specialization in preying upon diatoms, suggesting a specific ecological niche associated with diatom availability.

The RF-based model and ANCOM-BC emphasised four major groups of prokaryotic ASVs associated with seasons, primarily affiliated with Proteobacteria, Bacteroidota and Verrucomicrobiota (Figure 3B). Based on those results we can speculate that the RF-selected ASVs not only reflect their response to the general ongoing seasonal changes in environmental conditions (such as temperature or DOC), but that their abundance might be promoted by their higher specialisation towards different abiotic and biotic factors than the dominant species. This division into widespread generalists and less numerous specialists, as well as niche partitioning in bacterioplankton communities, is well supported by various research in both marine and freshwater ecosystems (Gómez-Consarnau et al., 2012; Liao et al., 2016; Mo et al., 2021). Interestingly, specialists in aquatic systems are frequently involved in the degradation of dissolved organic carbon (DOCp) from phytoplankton, which is produced by exudation or cell lysis (Sarmento et al., 2016). Combined with the fact that different types of phytoplankton promote the growth of different bacterial groups (Sarmento & Gasol, 2012), our results suggest that many of the observed intermittent occurrences of ASVs are due to such associations. The ASVs selected by RF were not only more diverse (even at the phylum level) than the dominants, but also belonged to groups previously reported to be involved in the degradation of certain organic compounds produced by phytoplankton. This is most evident in April, where the prolonged diatom bloom stimulated the growth of ASV117 (*Rhodobacter*), ASV113 (Verrucomicrobiae) and members of the Bacteroidota (ASV132, ASV142, ASV154), which are either specialised in the degradation of diatom DOCp or are generally associated with diatom blooms (Tada et al., 2012; Orellana et al., 2022). Furthermore, a similar assemblage of taxa (Verrucomicrobiae and Bacteroidota) was also present in June, although we cannot explicitly point to the source of the organic matter, which could be either from the RF-selected chlorophytes and raphidophytes, or other algae (Figure 3A). In the ‘late summer’ assemblage, we also observed taxa such as the genus *Fluviicola* (ASV170), which has been reported to be closely associated with blooms of primary producers (Eckert et al., 2012). Identification of a single ASV assigned to *Cyanobium*, Cyanobacteria (ASV139) during the summer months (June to September) corresponds to the expected cyanobacterial bloom typical for this period. However, due to the two-stage filtration approach (excluding particles larger than 12 μm) we weren’t able to detect larger, colonial species like *Microcystis* or *Dolichospermum* spp., which are typically observed during cyanobacterial blooms (Woodhouse et al., 2018).

Distinct seasonal patterns are not only more evident for eukaryotes than prokaryotes, but they are also more stable when facing short-term environmental fluctuations. In samples A1 from July, we noted appearance of ASVs associated with spring-related assemblage of bacterial ASVs. Probably primarily caused by a drop in the water temperature (from ∼23 °C in June to ∼20 °C in July) as a result of rainfalls, low air temperature which subsequently increased relative abundance of diatoms - ASV3 (up to 5%) that subsequently stimulated growth diatom DOCp specialists (Figure 2A and 3B, Supplementary Table S2). Such environmental change exerted a more pronounced influence on the prokaryotic community compared to the eukaryotic community (Jacobsen & Simonsen, 1993; Stockwell et al., 2020). This observation might be explained by the specificity of the sampling site – a shallow, sheltered bay, which resembles other shallow water bodies, such as ponds, which are more responsive to physicochemical changes like aforementioned temperature, rainfalls or solar radiation, because of their low buffering capacity within limited water column (Simon et al., 2015b). The rapid change in prokaryotic community composition could be furtherly explained by higher cell division rates of prokaryotes than eukaryotes, which enables rapid adjustment to environmental conditions (Logares et al., 2015). These findings also prove that the monthly sampling scheme is not sufficient to determine the factors influencing the changes in microbial communities, because their turnover occurs rather in days than weeks (Šimek et al., 2014).

### 4.4 Microbial community of the anoxic hypolimnion

The majority of research on hypolimnion ecology focused on deep freshwater lakes with oxygenated hypolimnion, leading to the identification of distinct microbial communities in these environments (Okazaki & Nakano, 2016; Mukherjee et al., 2017). However, anthropogenic eutrophication of lakes and climate change increases the number of lakes experiencing anoxic conditions in their hypolimnion during the summer months and it is reasonable to anticipate that such anoxic environments would also harbour unique microbial communities. Surprisingly, studies investigating protistan communities in anoxic hypolimnion have been relatively scarce thus far. A case in point is the Lake Roś, which exhibits anoxic conditions within its hypolimnion during the summer. This environmental characteristic is further reflected in the establishment of a distinct hypolimnetic microbial community (Figure 4A). We propose that the main driver of this microbial community is decomposition of organic matter by methanogens – a phenomenon well-documented in other lakes exhibiting similar conditions characterized by high redox potential during stratification (Shi et al., 2022; Steinsdóttir et al., 2022; Reis et al., 2022). While our study using the V4 16S rDNA marker was unable to detect the presence of Archaea due to primer incompatibility, the analysis of V9 eukaryotic 18S rDNA data, with its capacity to amplify small ribosomal subunits from all three domains, revealed a higher relative abundance of various archaeal lineages in B2 samples compared to other sites and depths, reaching up to 1%, including the methanogenic *Methanosarcina* (Supplementary Figure S2) (Choi & Park, 2020; Carr & Buan, 2022). Consequently, we conclude that methane may be mainly oxidized by *Methylobacter* (ASV30) that uses alternate electron acceptors under anoxic conditions (Hao et al., 2022). Recent reports, also suggest the strong syntrophy between *Methylobacter* and *Methylotenera* (ASV7), which was the dominant methylotrophic genus that accounted for up to 9.5% of relative abundance. Their relationship couples nitrogen and carbon (C1) cycles with the extensive use of nitrates as alternative electron acceptors, which helps transfer carbon from methane to other members of hypolimnetic food web, such as Bacteroidota (ASV9 and ASV29), or the methylotrophic *Methylophilus* (ASV8) (Van Grinsven et al., 2021). In addition to the observed impact of biomass influx from the upper layer of the lake on the microbial community structure within the hypolimnion, there are reports suggesting the significance of other compounds often found in high concentrations, such as iron and manganese, in shaping the community. High concentrations of iron were observed in samples from site B of Lake Roś and can be associated with the growth of specific bacterial taxa, such as *Candidatus* Omnitrophus which relies on iron for magnetosome biosynthesis (Kolinko et al., 2016).

Within this ecosystem, protists mainly play the role as bacterial grazers. These include the genus *Bodo* (ASV171) and a choanoflagellate (ASV261), which have previously been reported in various marine and brackish anoxic environments (Bernard et al., 2000; Stock et al., 2009). Notably, we also identified an unknown eukaryotic lineage represented by ASV200 (with a low sequence identity ∼84% to an unknown eukaryote), suggesting the potential for the discovery of novel eukaryotic lineages within such ecosystems. Our data show that a well-established hypolimnetic community is formed in August, whereas, during the months of June and July, eukaryotes were dominated by ASVs related to Fungi or stramenopiles (diatoms), suggesting a potentially significant influence from the influx of dead algae, while eukaryotic ASVs associated with anoxia were present during this period in low abundance. These findings suggest that sinking dead cells and organic particles are vital early contributors to eukaryotic and bacterial plankton community development in the hypolimnion as a source of organic matter. An open question remains regarding the persistence of hypolimnion-associated lineages during the biannual water column mixing in spring and autumn. Most likely their refuge is in the sediments, often inhabited by obligately anoxic benthic microbiomes (Bernhard et al., 2014; Gomaa et al., 2022).

### 4.5 Abiotic and biotic factors shaping microbial community

We conducted an analysis of various abiotic factors previously shown to influence microbial communities (Bock et al., 2020). Among these factors, only a subset was found to be pivotal in shaping the microbial communities of Lake Roś. Temperature, dissolved oxygen, and silicon concentration emerged as the primary drivers of both communities’ structures (Figure 6). While temperature and dissolved oxygen levels have been recognized as significant factors shaping microbial community composition in many studies (Liu et al., 2013) Oliverio et al., 2018; Boenigk et al., 2018; Mikhailov et al., 2022; Shang et al., 2022), the silicon concentration seems to be mainly related to temperate lakes, with spring and autumn mixing events (Panizzo et al., 2018; Kong et al., 2021). In Lake Roś, the silicon concentration reaches its maximum during the autumn mixing, followed by the diatom bloom in spring, leading to a decrease in silicon concentration due to its utilization by diatoms for building their cell walls. A similar trend has been observed in Lake Baikal (Mikhailov et al., 2022). Other factors, including trophic status and phosphorus and nitrogen concentrations, corresponded with the prokaryotic community structure, but did not show strong impact on the eukaryotic community. These results align with previous research in freshwater lakes, suggesting that changes in bacterial communities are more closely linked to physicochemical patterns compared to protist communities (Bock et al., 2020).

In our study we also unveiled the significant influence of zooplankton on the diversity of protists (Supplementary Figures S13 and S14), a critical component of freshwater food webs, as top-down regulators of larger protists (Lu & Weisse, 2022). This impact became evident on a global scale through the correlation between protistan ASVs and zooplankton cell counts, particularly in the case of the predatory omnivorous rotifer *Asplanchna,* and the cladoceran *Diaphanosoma brachyurum* with preference for smaller (<3 μm) particles (Chang et al., 2010; Nandini et al., 2021). Furthermore, at a local scale, this influence was evident in beta-diversity patterns. We observed an unusual clustering pattern among samples from May (Supplementary Figure S12), suggesting that zooplankton, through their grazing activity (top-down selection) during clear water phase, could dramatically alter the local composition of protists in a specific region within the lake. In such cases, this impact has the potential to disrupt the balance between various protist groups.

### 4.6 Climate change Impact on microbial communities in dimictic temperate lakes

Our data clearly shows the pivotal role of temperature and oxygen levels in shaping distinct planktonic communities (Figure 6). Rapid changes in these factors, driven by climate change, are expected to have increasingly profound effects on freshwater lake ecosystems, thereby impacting biodiversity and ecosystem functions. Notably, surface water temperatures of freshwater ecosystems are rising at an accelerated rate compared to air and ocean temperatures (O’Reilly et al., 2015; Dokulil et al., 2021). Recent studies have demonstrated that warming predominantly leads to a decrease in freshwater plankton diversity (Rasconi et al., 2017; Urrutia-Cordero et al., 2017; Bergkemper et al., 2018; Verbeek et al., 2018; Da Silva et al., 2019; Celewicz & Gołdyn, 2021). However, the effects are multifaceted, often resulting in shifts in community structures, particularly towards green algae dominance (Rasconi et al., 2017; Yu et al., 2018; Zhang et al., 2021; Beng et al., 2023). Considering the complexity of these changes, it is essential to assess diversity at an appropriate taxonomic level, since the shifts might not be evident at higher taxonomic levels. Our study illustrates that the abundance of specific ASVs can exhibit dynamic changes over time, even when the abundance of the broader taxonomic groups they belong to remains relatively stable (Figures 2A and 2B).

Additionally, climate change is driving a decline in dissolved oxygen levels in aquatic ecosystems, affecting lakes, coastal zones, and open oceans globally (Breitburg et al., 2018; Schmidtko et al., 2017; Limburg et al., 2020; Jane et al., 2021). Large-scale analyses reveal that the majority of lakes (over 70%) are experiencing increases in oxygen-depleted water (Jane et al., 2023). This trend is significant since reduced dissolved oxygen concentrations can be observed during late summer periods due to changes in stratification characteristics (Jane et al., 2023), including earlier onset of seasonal stratification and less frequent mixing events (Woolway & Merchant, 2019). We have identified a distinct hypolimnetic community (Figure 4, Supplementary Table S8) that thrives in oxygen-depleted waters. This community is primarily composed of kinetoplastids (ASV171), choanoflagellates (ASV261), an amoebozoan (ASV116), and ASV200 from a novel protist group. While their role in the lake’s ecosystem is not as well understood as that of epilimnetic plankton, the increase in hypoxic zones highlights the critical need to explore the taxonomic and functional diversity of this community.

## 5. Conclusions

In our metabarcoding study of a typical temperate dimictic lake, we gained insights into the taxonomic composition and community structure of microbial eukaryotes and prokaryotes during the ice-free period at the unprecedented level of single ASVs. Leveraging random forest and ANCOM-BC analyses, we identified ecologically functional clusters of eukaryotic and prokaryotic ASVs that are specifically associated with different seasons. In particular, certain protist ASVs showed robust seasonal changes and played a crucial role in altering community structure between seasons. These changes were mainly associated with consumer groups such as cercozoans, and parasitic taxa such as Pseudofungi and Chytridiomycota. In contrast, the generalist ASVs, at least during the ice-free season, were mainly phototrophic and mixotrophic organisms, such as Cryptophyta, and predators, such as Katablepharidophyta. The prokaryotic diversity could also be divided into generalists (such as Actinobacteria) and specialists, which are a diversified group of taxa that are most likely involved in recycling organic matter, such as DOCp, abundant at certain time points.

Besides seasonal communities, significant differences were also observed between microbial communities in the epilimnion and hypolimnion, with key hypolimnion-specific taxa identified, including Choanoflagellata, Amoebozoa (Lobosa), Discoba (Kinetoplastida), and a putative novel lineage (ASV200). These taxa likely feed on a prokaryotic community driven by organisms involved in C1 cycle, such as methanogens and methanotrophs. The observed differential seasonal patterns in protistan and prokaryotic communities align with their distinct responses to environmental factors. Eukaryotes exhibited different responses compared to prokaryotes, particularly to the three main factors of temperature, oxygen, and silicon concentration. While these factors affected both groups, other environmental variables primarily influenced bacterial communities (bottom-up regulation). In contrast, zooplankton composition and abundance exerted a more pronounced top-down influence on the eukaryotic community compared to the prokaryotic community. In summary, our research provides significant insights into the seasonal dynamics of prokaryotic and eukaryotic communities in a temperate dimictic lake and reveals the ASVs that drive these communities using metabarcoding and machine learning methods.

## Acknowledgements

This work was supported by the European Molecular Biology Organization and Polish Ministry of Education and Ministry of Education and Science, Poland [EMBO Installation Grant 4150 to AK] and National Science Centre, Poland [OPUS grant 2020/37/B/NZ8/01456 to AK]. Sampling was carried out with the equipment of the Hydrobiological Field Station of the Faculty of Biology. We would like to thank everyone who helped us with the sampling, especially the head of the station Mirosław Ślusarczyk, and Maria Turłaj.

## Autor Contributions

AK designed the work and organized the sampling. MK, AB, PH, KM and AK collected and processed the samples. AB and MK purified DNA and carried out PCR reactions for metabarcoding analysis. MK carried out bioinformatic analysis of amplicon sequences and statistical analyses. MK wrote the first draft of the manuscript. MK and AK wrote the final version of the manuscript with input of other co-authors. All authors read, critically commented and approved the final manuscript.

