## Supplementary Figures and Data for "Seasonal dynamics and drivers of microbial communities in a temperate dimictic lake: insights from metabarcoding and machine learning"

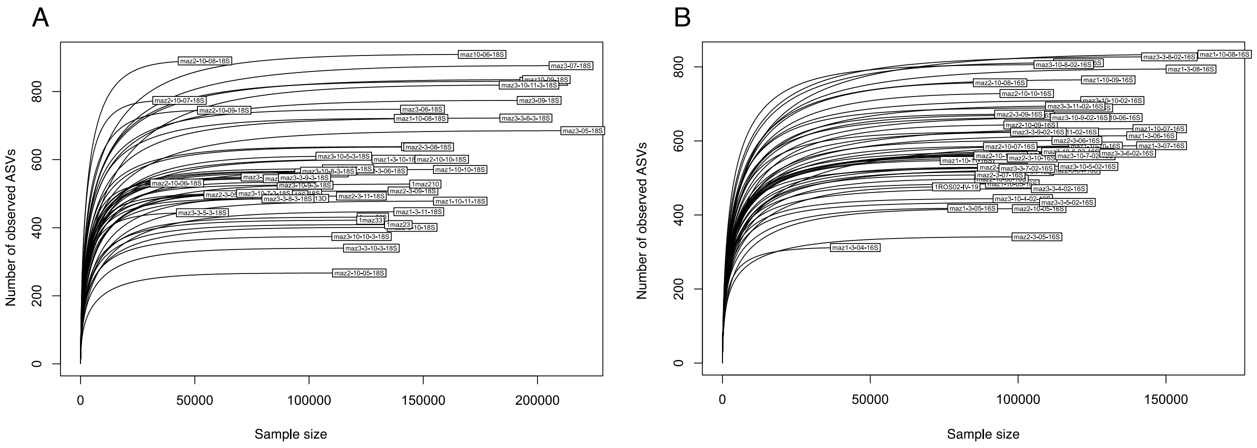


**Supplementary Figure S1.** Rarefaction curves for (A) 18S V9 and (B) 16S V4 datasets based on the number of reads (sample size) and the number of ASVs in each sample. Each line represents one sample.

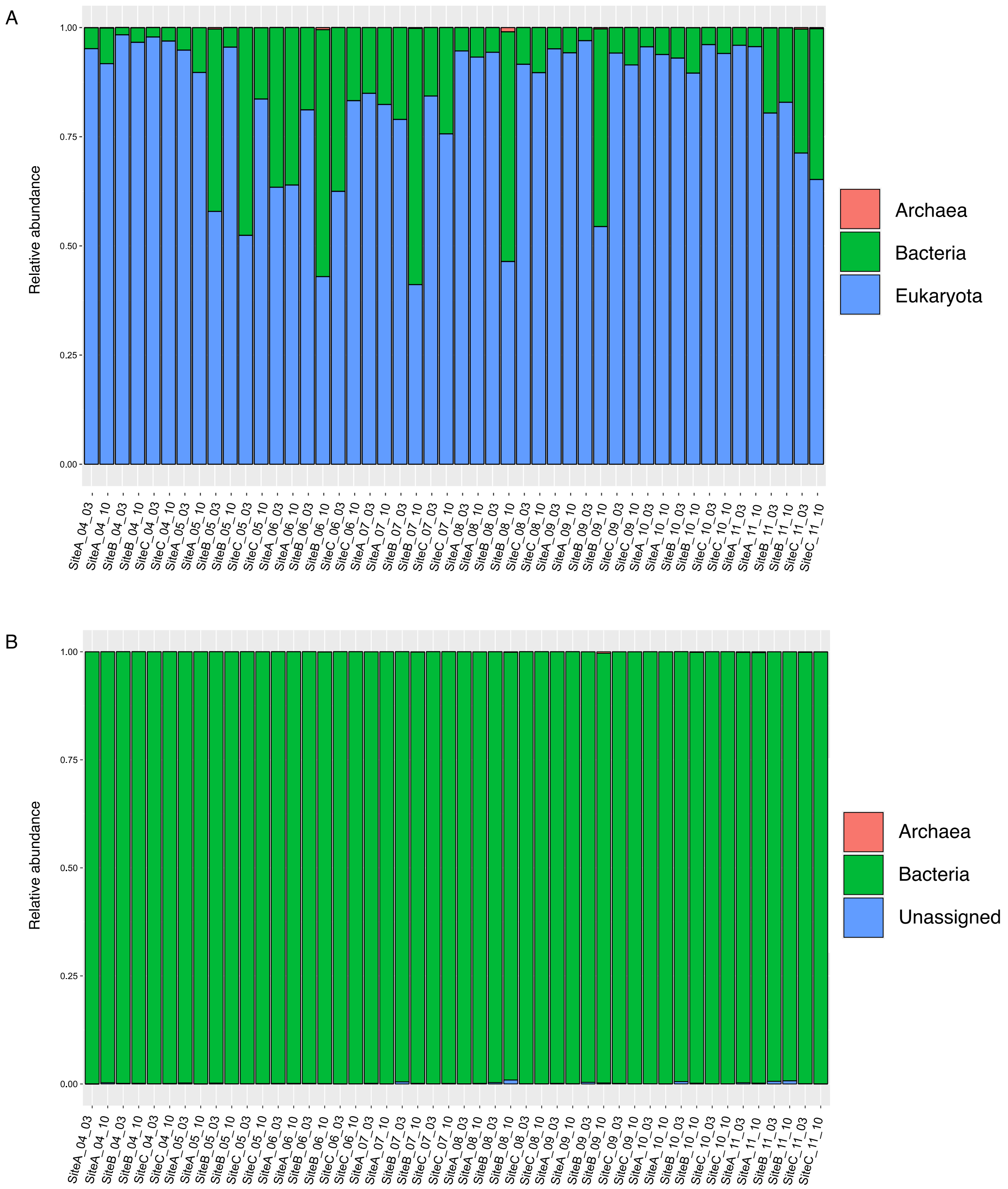


**Supplementary Figure S2.** Relative abundance at domain level across samples for (A) 18S V9 rDNA and (B) 16S V4 rDNA. Representation of the proportion of incorrectly amplified bacterial and archaeal ASVs (A) and the low presence of archaeal ASVs in the prokaryotic data set (B).

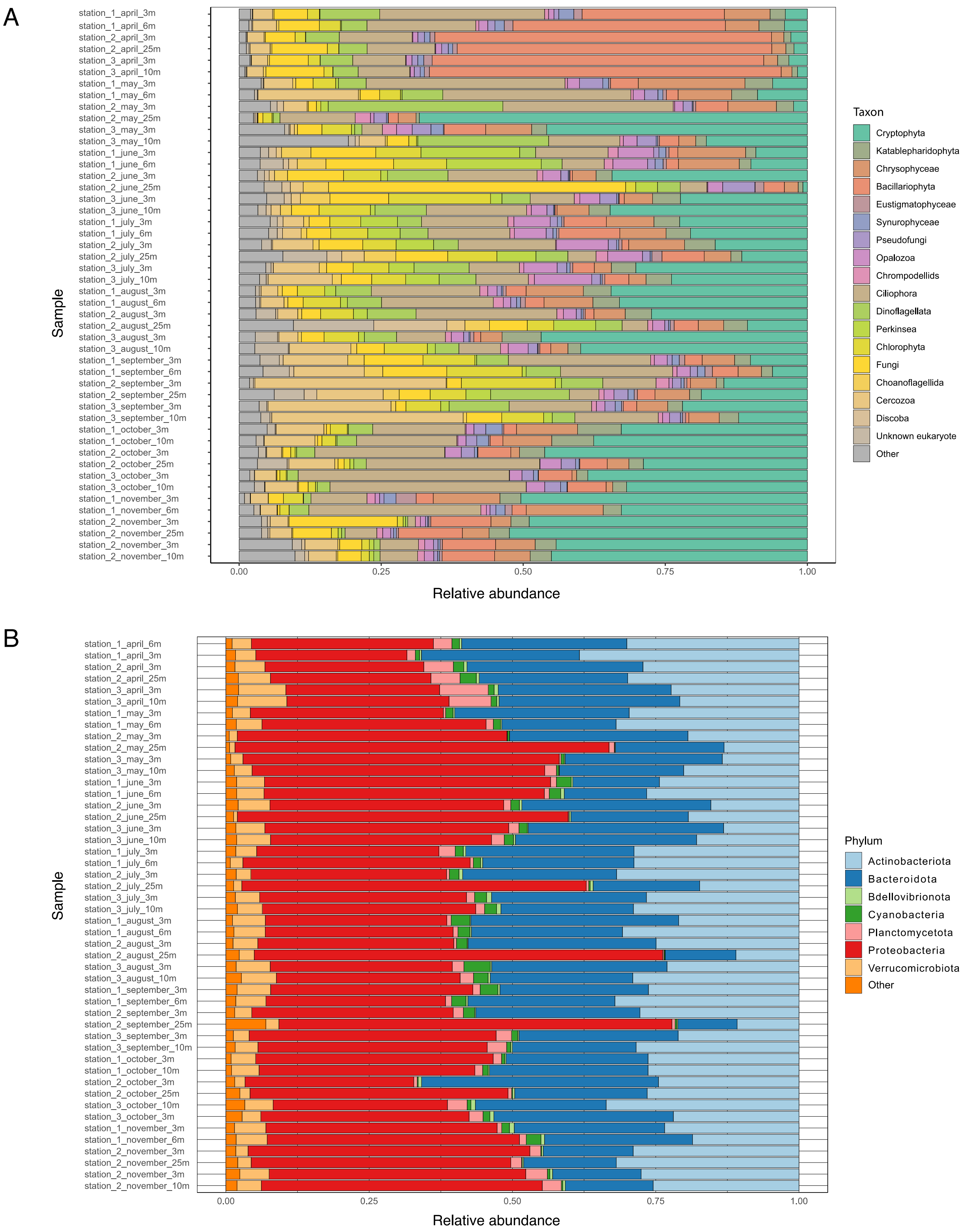


**Supplementary Figure 3.** Relative abundance across samples (A) at the 'Class' level as defined by the pr2 database for protist 18S V9 rDNA ASVs, and (B) at the 'phylum' level as defined by the SILVA database for prokaryotic 16S V4 ASVs.

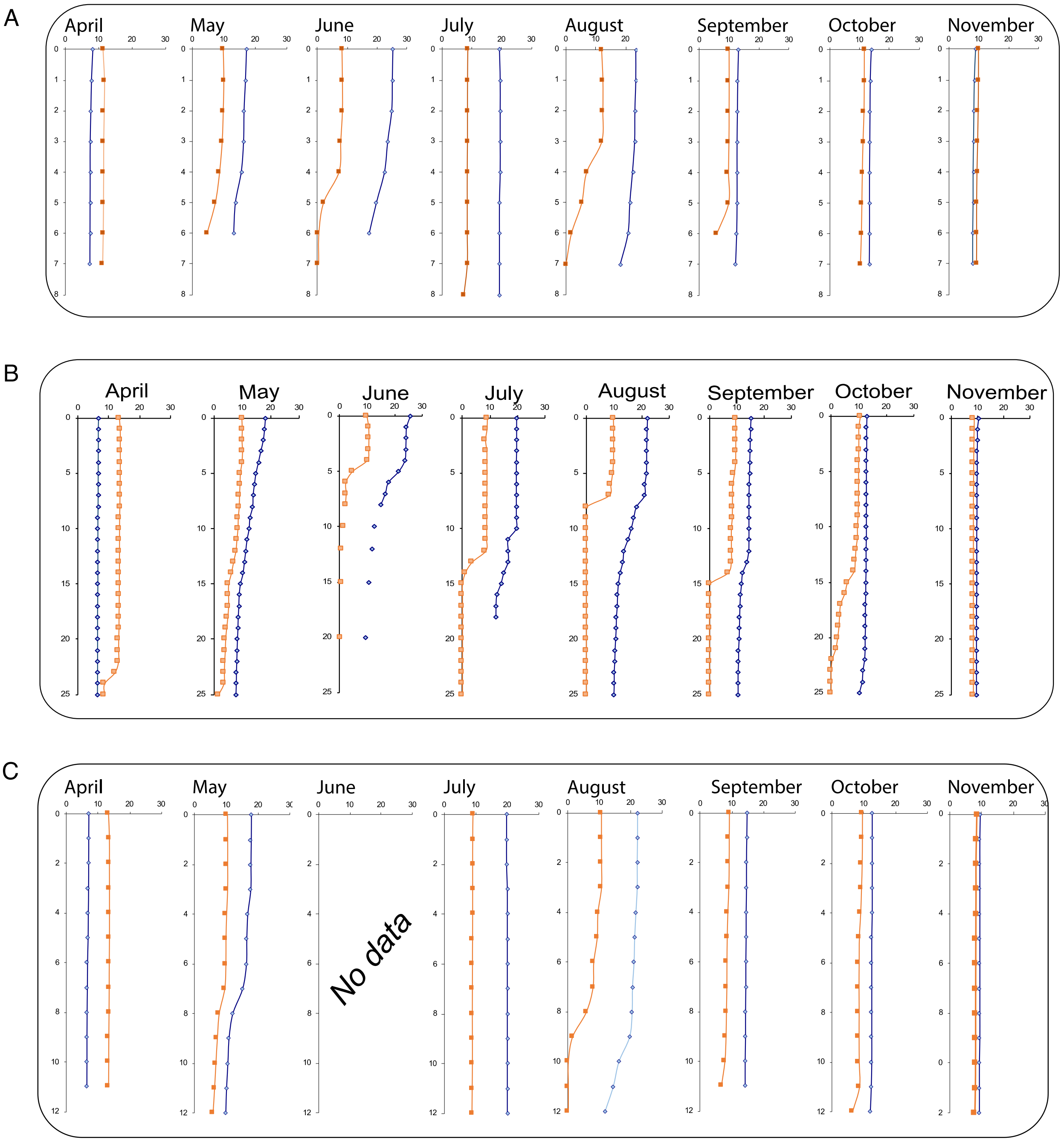


**Supplementary Figure S4.** Vertical oxygen (marked orange) and temperature (marked blue) profiles for (A) Site A (8 m), (B) Site B (25 m), and (C) Site C (12 m). The missing data in June at site C was caused by technical problems with the multiparametric probe.

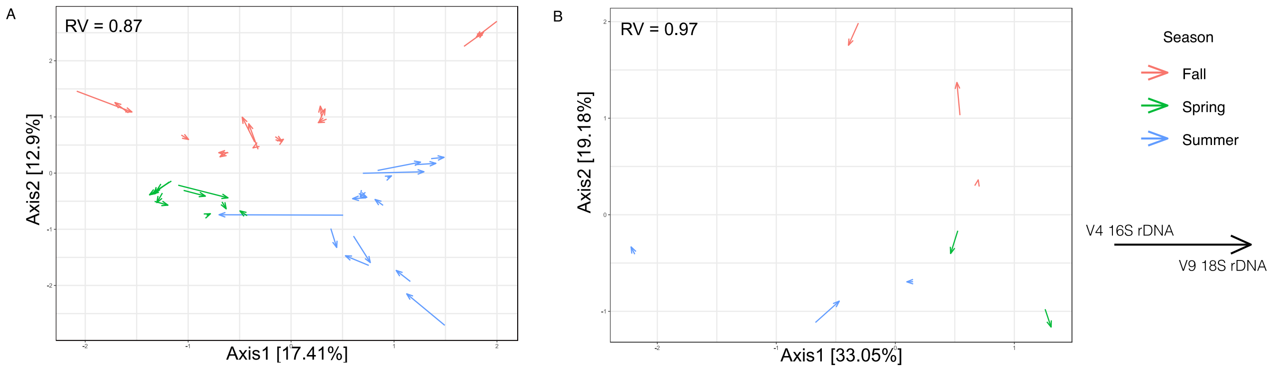


**Supplementary Figure S5**. PCA-based co-inertia analysis (A) for all samples except 25 m (A1, A2, B1, C1 and C2) and (B) for the sample at 25 m (B2).

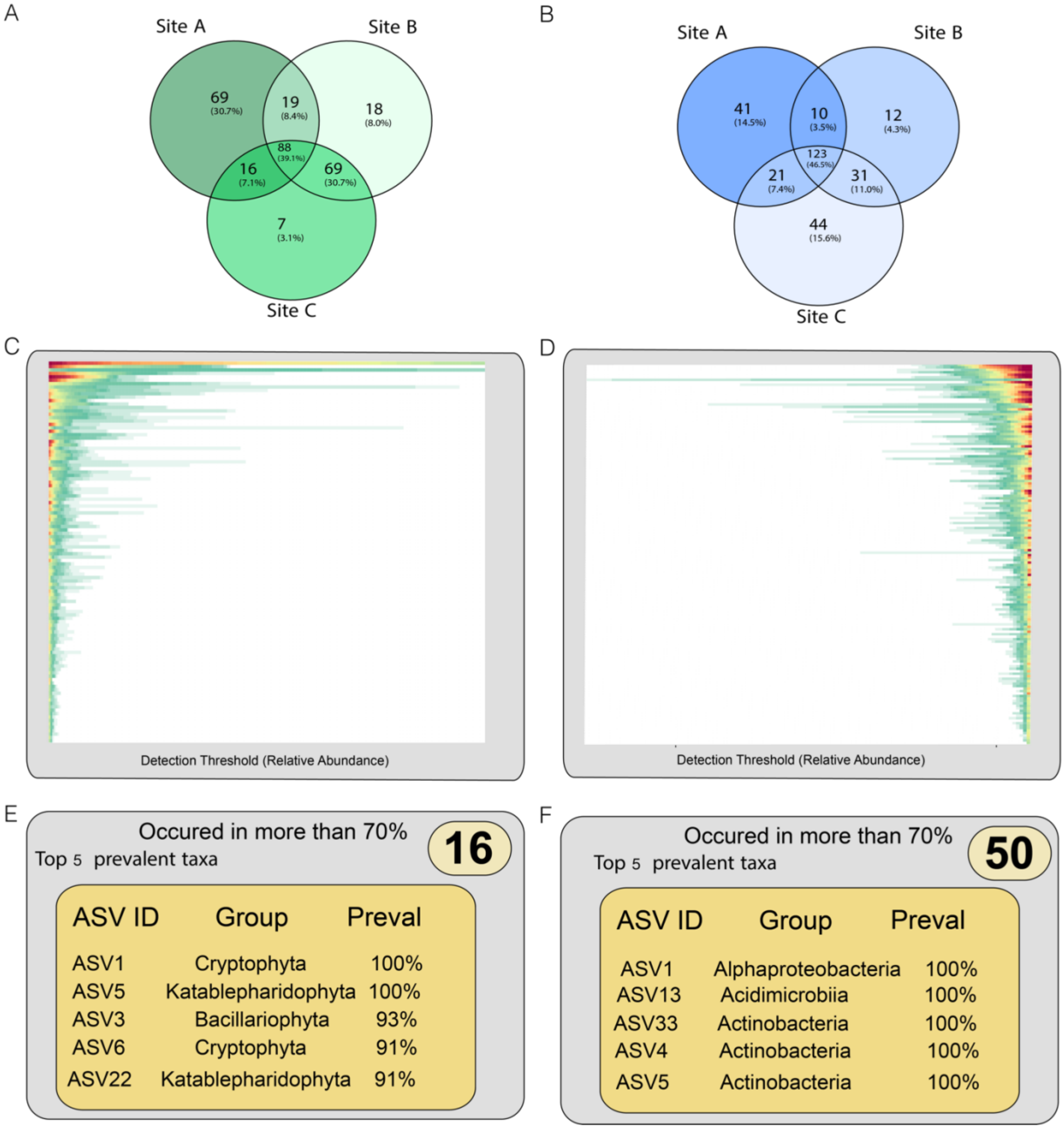


**Supplementary Figure S6.** Venn diagrams showing the occurrence of (A) eukaryotic and (B) prokaryotic ASVs at sites A, B and C for the entire sampling season. Analysis of the prevalence of (C) eukaryotic and (D) prokaryotic ASVs in samples A1, A2, B1, C1, C2 across the sampling seasons (all samples except 25 m). The most prevalent (E) eukaryotic and (F) prokaryotic ASVs are present in more than 70% of the samples at least in relative abundance 0.001.

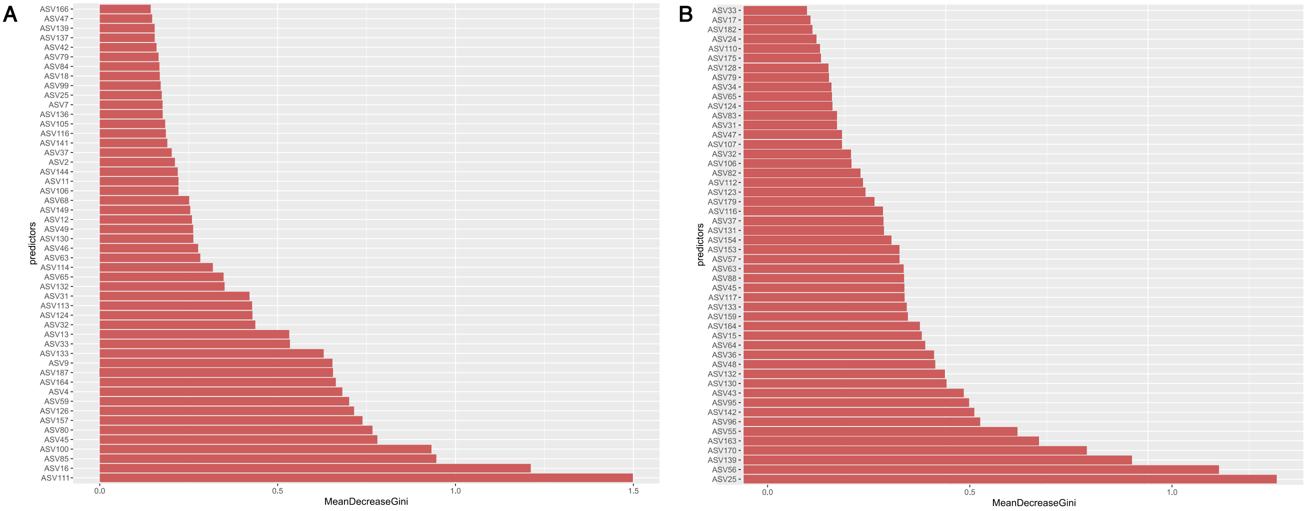


**Supplementary Figure S7.** Mean Gini indices of (A) eukaryotic and (B) and prokaryotic predictors selected by random forest models.

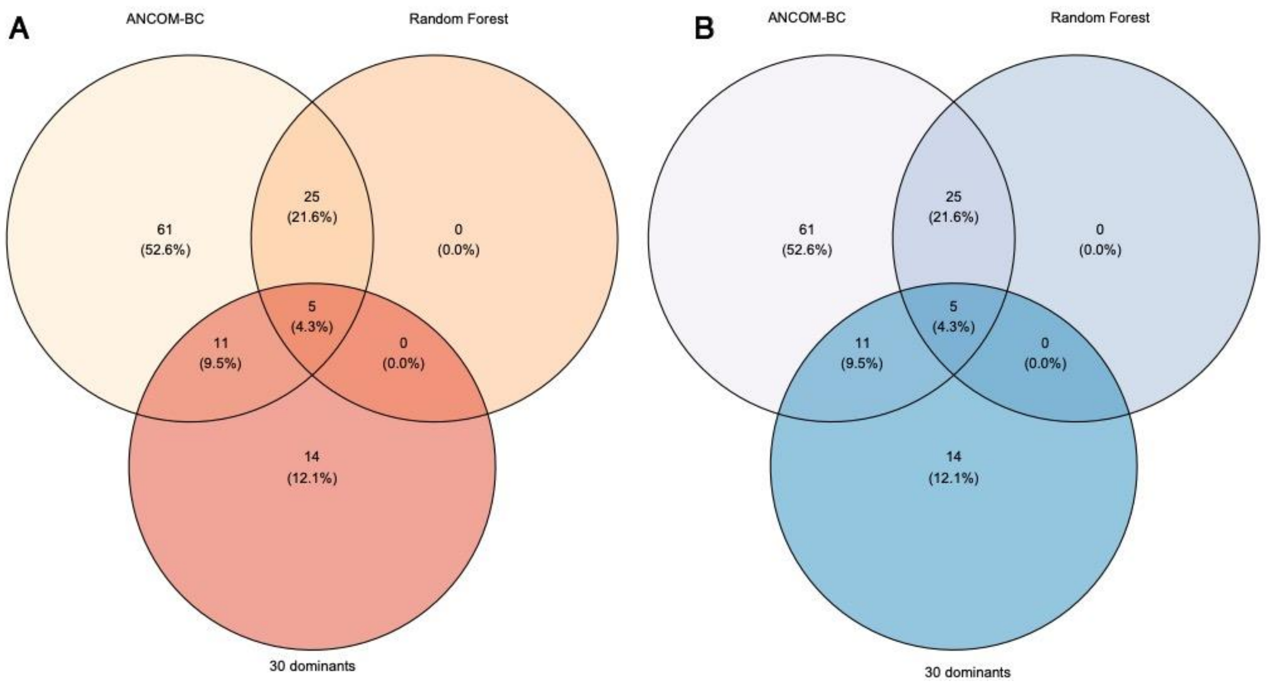


**Supplementary Figure S8.** Venn diagrams of (A) eukaryotic 18S V9 rDNA and (B) prokaryotic 16S V4 rDNA ASVs selected by Ancom-BC and Random-Forest methods compared to 30 dominant ASVs.

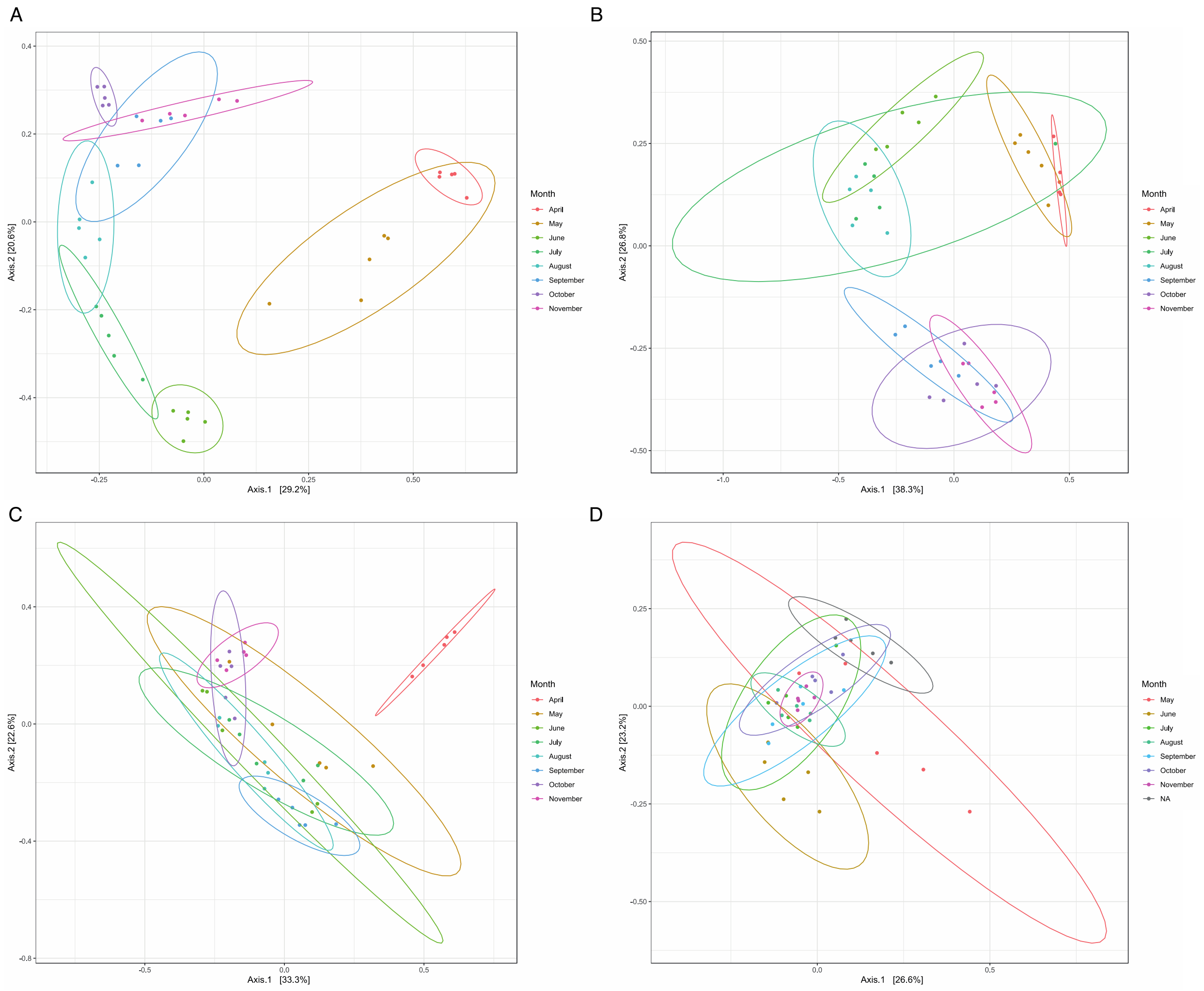


**Supplementary Figure 9.** Bray-Curtis MDS plots based on (A) eukaryotic and (B) prokaryotic ASVs selected by Random Forest models compared with Bray MDS plots based on 30 dominant ASVs for (C) eukaryotes and (D) prokaryotes. The colors represent months.

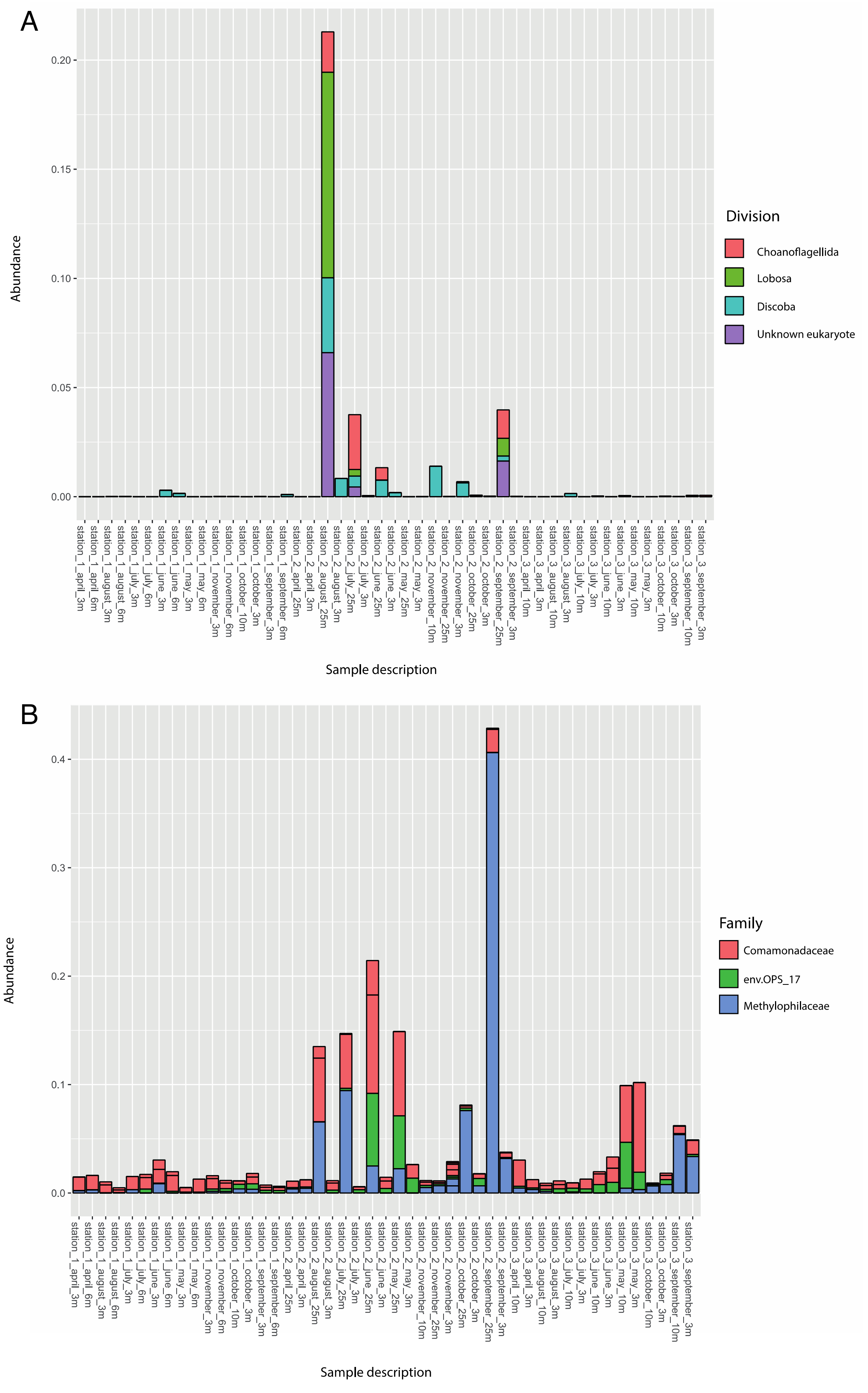


**Supplementary Figure S10.** Relative abundance of (A) eukaryotic and (B) prokaryotic ASVs associated with the hypolimnion in all samples. The colours refer to the taxonomic affiliation of the ASVs.

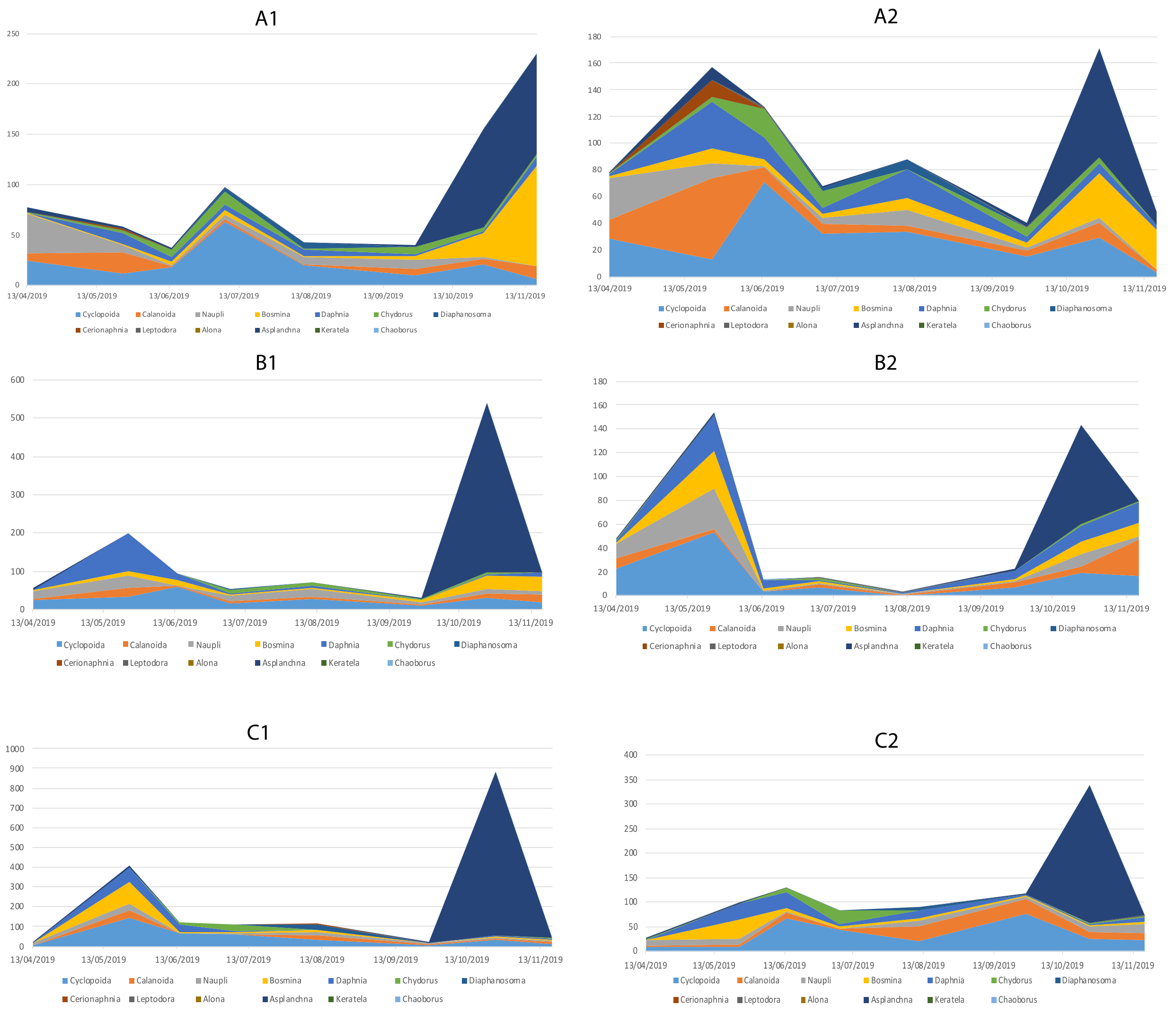


**Supplementary Figure S11.** Microscopically determined absolute counts of groups of zooplankton (number of specimens per liter) for each site and depth across sampling season.

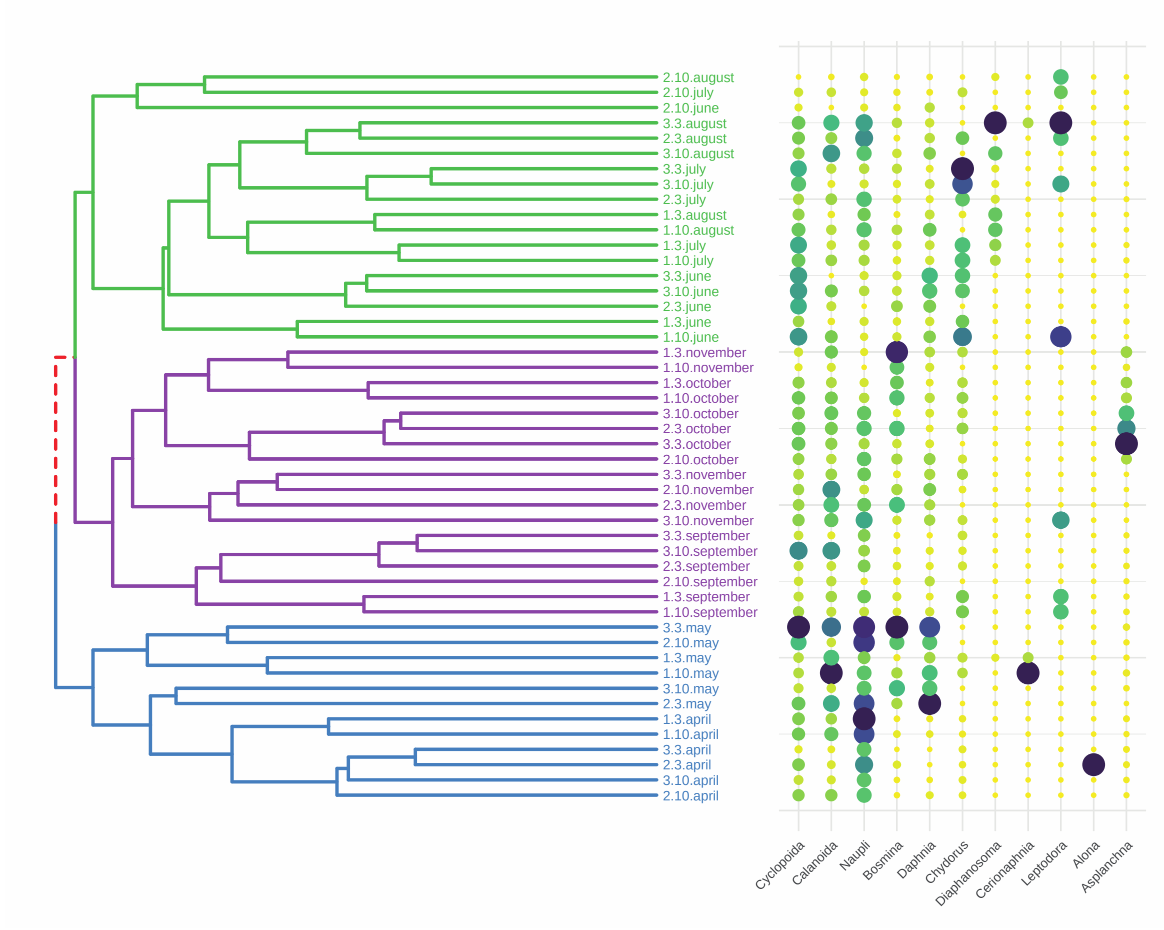


**Supplementary Figure S12.** Dendrogram of eukaryotic samples (18 V9 rDNA) based on the Unweighted UniFrac metric ("complete" clustering method) compared to the absolute counts of the different zooplankton groups (number of specimens per liter).

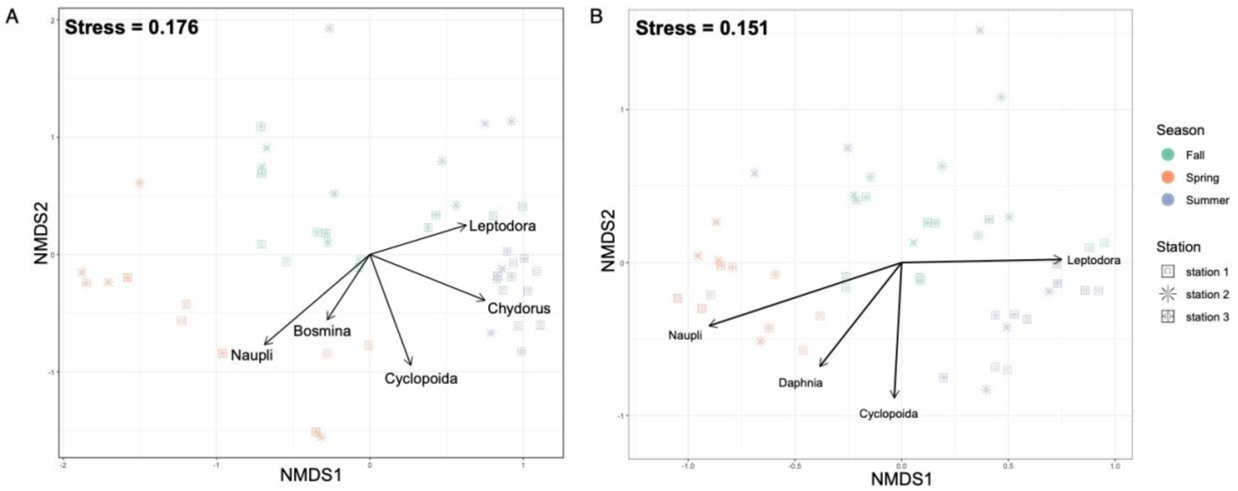


**Supplementary Figure S13.** Sample composition of (A) eukaryotic and (B) prokaryotic datasets based on NMDS analysis with envfit (vegan package) fitted zooplankton absolute counts ($p < 0.05$). The colors refer to the seasons, the shapes to the sampling sites.

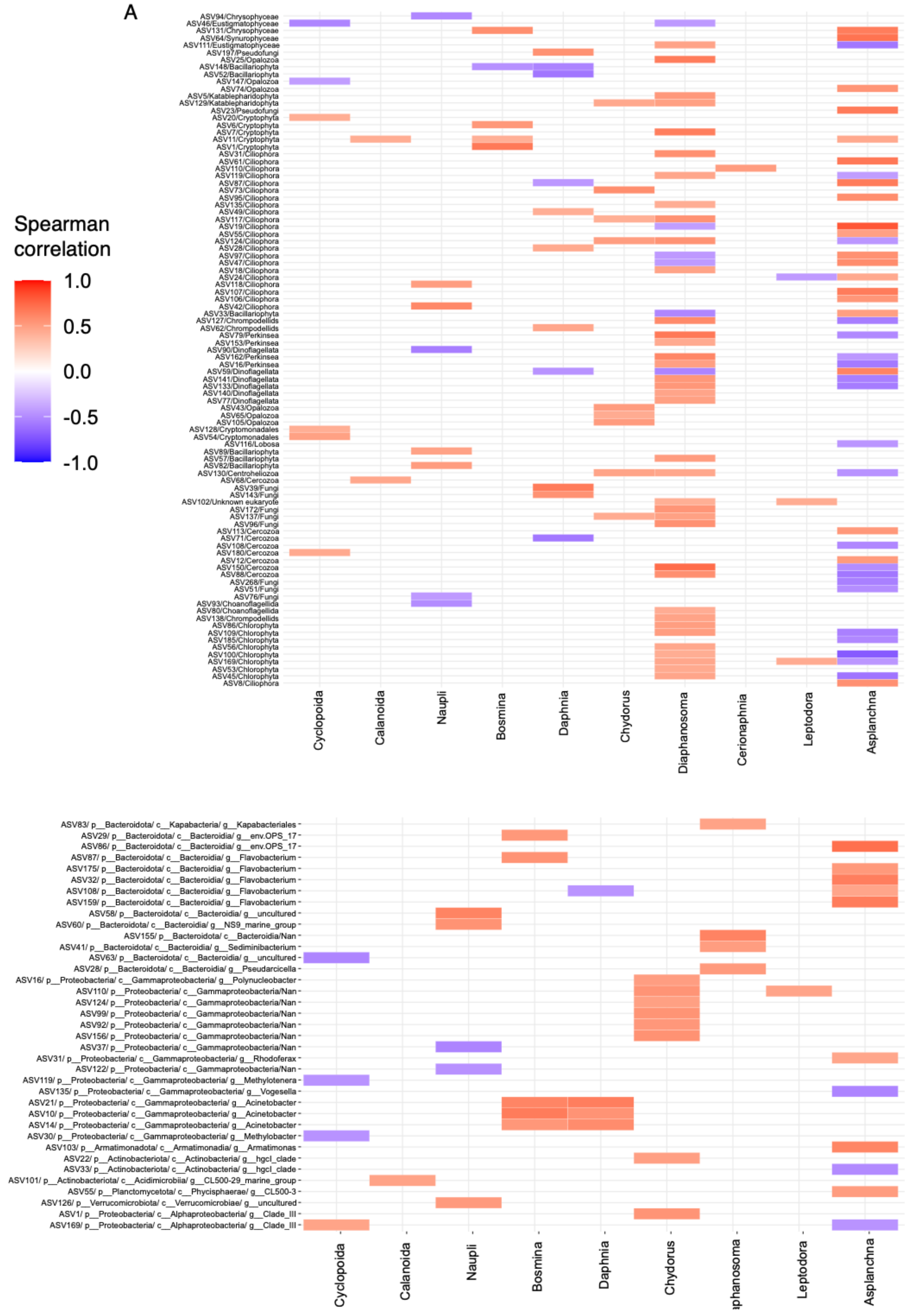


**Supplementary Figure S14.** Spearman correlogram between ASVs and absolute counts of zooplankton groups ($p < 0.05$) for (A) eukaryotic and (B) prokaryotic ASVs with a relative abundance of more than 0.001. The colors refer to the character of correlation (positive – red, blue – negative) and the intensity to the R value.

**Supplementary Data 1**

Presents results of the analysis of the eukaryotic part of the data analysis (V9 18S rDNA dataset).

### Alpha diversity

alpha_stats.shannon.anova <- aov(Shannon ~ seqsDepth + Season + Month + Station + Depth..m. * Month, dframe_alpha_to_stats)
summary(alpha_stats.shannon.anova)

```
   Df Sum Sq Mean Sq F value   Pr(>F)
Season         2  6.353   3.176  34.896 1.48e-08 ***
Month          5  4.956   0.991  10.889 4.86e-06 ***
Station        2  2.584   1.292  14.192 4.60e-05 ***
Depth..m.       1  0.141   0.141   1.553 0.222307
Month:Depth..m.  7  4.150   0.593   6.513 0.000103 ***
Residuals      30  2.731   0.091
---
Signif. codes:  0 ‘***’ 0.001 ‘**’ 0.01 ‘*’ 0.05 ‘.’ 0.1 ‘ ’ 1
```

### Beta diversity

Adonis analysis of selected metrics with three factors: Station (A/B/C), Season (spring/summer/autumn), Month (April, March, April, May, June, July, August, September, November)

```
adonis2(dist_wunifrac_otus_euk ~ Station, data = table_metadata)
adonis2(dist_wunifrac_otus_euk ~ Season, data = table_metadata)
adonis2(dist_wunifrac_otus_euk ~ Month, data = table_metadata)
```

Permutation test for adonis under reduced model
Terms added sequentially (first to last)
Permutation: free
Number of permutations: 999

**~Station**

```
adonis2(formula = dist_wunifrac_otus_euk ~ Station, data = table_metadata)
       Df SumOfSqs      R2      F Pr(>F)
Station   2 0.005097 0.03919 0.9178   0.53
Residual 45 0.124941 0.96081
Total    47 0.130037 1.00000
```

**~Season**

```
adonis2(formula = dist_wunifrac_otus_euk ~ Season, data = table_metadata)
       Df SumOfSqs      R2      F Pr(>F)
Season    2 0.032086 0.24675 7.3704  0.001 ***
Residual 45 0.097951 0.75325
Total    47 0.130037 1.00000
---
Signif. codes:  0 ‘***’ 0.001 ‘**’ 0.01 ‘*’ 0.05 ‘.’ 0.1 ‘ ’ 1
```

Permutation test for adonis under reduced model
Terms added sequentially (first to last)
Permutation: free
Number of permutations: 999

**~Month**

```
adonis2(formula = dist_wunifrac_otus_euk ~ Month, data = table_metadata)
        Df SumOfSqs      R2      F Pr(>F)
Month     7 0.080153 0.61638 9.1815  0.001 ***
Residual 40 0.049885 0.38362
Total    47 0.130037 1.00000
---
Signif. codes:  0 '***' 0.001 '**' 0.01 '*' 0.05 '.' 0.1 ' ' 1
#Beta-dispersion vs Season
disp_Season_unwuni = betadisper(dist_ununifrac_otus_euk, table_metadata$Season)

#Beta-dispersion vs Month
disp_Month_unwuni = betadisper(dist_ununifrac_otus_euk, table_metadata$Month)

#Beta-dispersion vs Station
disp_Station_unwuni = betadisper(dist_ununifrac_otus_euk, table_metadata$Station)

permutest(disp_Season_unwuni, pairwise=TRUE, permutations=1000)

permutest(disp_Month_unwuni, pairwise=TRUE, permutations=1000)

permutest(disp_Station_unwuni, pairwise=TRUE, permutations=1000)
```

**~Season**

```
Permutation test for homogeneity of multivariate dispersions
Permutation: free
Number of permutations: 1000

Response: Distances
          Df   Sum Sq   Mean Sq      F N.Perm Pr(>F)
Groups     2 0.004619 0.0023097 0.4523   1000 0.6384
Residuals 45 0.229783 0.0051063

Pairwise comparisons:
(Observed p-value below diagonal, permuted p-value above diagonal)
          Fall  Spring Summer
Fall           0.84815 0.4126
Spring 0.83153         0.4476
Summer 0.41615 0.45613
```

**~Month**

```
Permutation test for homogeneity of multivariate dispersions
Permutation: free
Number of permutations: 1000

Response: Distances
          Df  Sum Sq   Mean Sq      F N.Perm Pr(>F)
Groups     7 0.08490 0.0121284 1.4929   1000 0.1578
Residuals 40 0.32496 0.0081239

Pairwise comparisons:
(Observed p-value below diagonal, permuted p-value above diagonal)
              April    August      July      June       May  November   October September
April               0.3946054 0.5524476 0.2047952 0.0059940 0.0609391 0.6173826    0.4665
August    0.4067890           0.7922078 0.5824176 0.0899101 0.3976024 0.6633367    0.8671
July      0.5476888 0.8187021           0.4615385 0.0419580 0.2587413 0.8891109    0.9251
June      0.2028824 0.6042936 0.4660778           0.2877123 0.9310689 0.3456543    0.4585
May       0.0054157 0.0754562 0.0415447 0.3020317           0.0589411 0.0169830    0.0300
November  0.0452068 0.4057670 0.2555822 0.9327188 0.0659724           0.1578422    0.2747
October   0.6309538 0.6994042 0.8819215 0.3758099 0.0203884 0.1556375              0.8062
September 0.4608883 0.8831154 0.9241855 0.4992861 0.0368918 0.2641032 0.7968145
```

**~Station**

Permutation test for homogeneity of multivariate dispersions
Permutation: free
Number of permutations: 1000

Response: Distances

| | Df | Sum Sq | Mean Sq | F | N.Perm | Pr(>F) | |
|---|---|---|---|---|---|---|---|
| Groups | 2 | 0.013654 | 0.0068268 | 3.2729 | 1000 | 0.04895 | * |
| Residuals | 45 | 0.093864 | 0.0020859 | | | | |

---
Signif. codes: 0 ‘***’ 0.001 ‘**’ 0.01 ‘*’ 0.05 ‘.’ 0.1 ‘ ’ 1

Pairwise comparisons:
(Observed p-value below diagonal, permuted p-value above diagonal)

| | station 1 | station 2 | station 3 |
|---|---|---|---|
| station 1 | | 0.011988 | 0.3307 |
| station 2 | 0.007306 | | 0.1568 |
| station 3 | 0.321690 | 0.156293 | |

**Supplementary Data 2**

Supplementary materials have been divided into two sections. Section 1 presents results of the analysis of the eukaryotic part of the data analysis (V9 18S rDNA dataset), whereas Section 2 presents the prokaryotic part of the analysis (V4 16S rDNA dataset).

### Alpha Diversity

```
alpha_stats.shannon.anova <- aov(Shannon ~ seqsDepth + Season + Month + Station + Depth..m.*Month, dframe_alpha_to_stats)
summary(alpha_stats.shannon.anova)
```

```
 Df Sum Sq Mean Sq F value   Pr(>F)
Season          2 0.4537  0.2268   8.621  0.00134 **
Month           4 0.6687  0.1672   6.353  0.00105 **
Station         2 0.9477  0.4738  18.008 1.24e-05 ***
Depth..m.       1 0.0747  0.0747   2.838  0.10404
Month:Depth..m.  6 0.7906  0.1318   5.008  0.00157 **
Residuals      26 0.6841  0.0263
---
Signif. codes:  0 '***' 0.001 '**' 0.01 '*' 0.05 '.' 0.1 ' ' 1
```

### Beta Diversity

```
adonis2(prok_dist_ununifrac_otus ~ Station, data = table_metadata)
adonis2(prok_dist_ununifrac_otus ~ Season, data = table_metadata)
adonis2(prok_dist_ununifrac_otus ~ Month, data = table_metadata)
```

Permutation test for adonis under reduced model
Terms added sequentially (first to last)
Permutation: free
Number of permutations: 999

**~Station**

```
adonis2(formula = prok_dist_ununifrac_otus ~ Station, data = table_metadata)
         Df SumOfSqs      R2      F Pr(>F)
Station   2   1.4444 0.09846 2.4573  0.001 ***
Residual 45  13.2252 0.90154
Total    47  14.6696 1.00000
---
Signif. codes:  0 '***' 0.001 '**' 0.01 '*' 0.05 '.' 0.1 ' ' 1
```

Permutation test for adonis under reduced model
Terms added sequentially (first to last)
Permutation: free
Number of permutations: 999

**~Season**

```
adonis2(formula = prok_dist_ununifrac_otus ~ Season, data = table_metadata)
         Df SumOfSqs      R2      F Pr(>F)
Season    2   2.5118 0.17122 4.6484  0.001 ***
Residual 45  12.1578 0.82878
Total    47  14.6696 1.00000
---
Signif. codes:  0 '***' 0.001 '**' 0.01 '*' 0.05 '.' 0.1 ' ' 1
```

Permutation test for adonis under reduced model
Terms added sequentially (first to last)
Permutation: free
Number of permutations: 999

**~Month**

```
adonis2(formula = prok_dist_ununifrac_otus ~ Month, data = table_metadata)
         Df SumOfSqs      R2      F Pr(>F)
Month     7   5.3884 0.36732 3.3175  0.001 ***
Residual 40   9.2812 0.63268
Total    47  14.6696 1.00000
---
Signif. codes:  0 ‘***’ 0.001 ‘**’ 0.01 ‘*’ 0.05 ‘.’ 0.1 ‘ ’ 1
```

```
#Beta-dispersion vs Season
disp_Season_unwuni = betadisper(prok_dist_ununifrac_otus, table_metadata$Season)

#Beta-dispersion vs Month
disp_Month_unwuni = betadisper(prok_dist_ununifrac_otus, table_metadata$Month)

#Beta-dispersion vs Station
disp_Station_unwuni = betadisper(prok_dist_ununifrac_otus, table_metadata$Station)

# Now we can test the significance with using permutation
set.seed(27)
permutest(disp_Season_unwuni, pairwise=TRUE, permutations=1000)

permutest(disp_Month_unwuni, pairwise=TRUE, permutations=1000)

permutest(disp_Station_unwuni, pairwise=TRUE, permutations=1000)
```

```
Permutation test for homogeneity of multivariate dispersions
Permutation: free
Number of permutations: 1000
```

**~Month**

```
Response: Distances
          Df   Sum Sq   Mean Sq      F N.Perm Pr(>F)
Groups     2 0.004619 0.0023097 0.4523   1000 0.6414
Residuals 45 0.229783 0.0051063

Pairwise comparisons:
(Observed p-value below diagonal, permuted p-value above diagonal)
          Fall  Spring Summer
Fall           0.83417 0.4156
Spring 0.83153         0.4486
Summer 0.41615 0.45613

Permutation test for homogeneity of multivariate dispersions
Permutation: free
Number of permutations: 1000

Response: Distances
          Df  Sum Sq   Mean Sq      F N.Perm Pr(>F)
Groups     7 0.08490 0.0121284 1.4929   1000 0.1888
Residuals 40 0.32496 0.0081239

Pairwise comparisons:
(Observed p-value below diagonal, permuted p-value above diagonal)
              April    August      July      June       May  November   October September
April               0.4025974 0.5374625 0.2197802 0.0089910 0.0519481 0.6243756    0.4406
August    0.4067890           0.8131868 0.5564436 0.0659341 0.3966034 0.6873127    0.8851
July      0.5476888 0.8187021           0.4465534 0.0369630 0.2347652 0.8531469    0.9291
June      0.2028824 0.6042936 0.4660778           0.2897103 0.9390609 0.3586414    0.4865
May       0.0054157 0.0754562 0.0415447 0.3020317           0.0629371 0.0169830    0.0310
November  0.0452068 0.4057670 0.2555822 0.9327188 0.0659724           0.1438561    0.2827
October   0.6309538 0.6994042 0.8819215 0.3758099 0.0203884 0.1556375              0.7962
September 0.4608883 0.8831154 0.9241855 0.4992861 0.0368918 0.2641032 0.7968145
```

Permutation test for homogeneity of multivariate dispersions
Permutation: free
Number of permutations: 1000

Response: Distances

| | Df | Sum Sq | Mean Sq | F | N.Perm | Pr(>F) | |
|---|---|---|---|---|---|---|---|
| Groups | 2 | 0.013654 | 0.0068268 | 3.2729 | 1000 | 0.04895 | * |
| Residuals | 45 | 0.093864 | 0.0020859 | | | | |

---
Signif. codes: 0 ‘***’ 0.001 ‘**’ 0.01 ‘*’ 0.05 ‘.’ 0.1 ‘ ’ 1

Pairwise comparisons:
(Observed p-value below diagonal, permuted p-value above diagonal)

| | station 1 | station 2 | station 3 |
|---|---|---|---|
| station 1 | | 0.007992 | 0.3347 |
| station 2 | 0.007306 | | 0.1758 |
| station 3 | 0.321690 | 0.156293 | |

**Supplementary Data 3**

Synchrony RV.rtest results

All samples (epi- and mesolimion) except 25 meters site B2

```
Monte-Carlo test
Call: RV.rtest(df1 = new_df, df2 = new_df2, nrepet = 99)

Observation: 0.8682183

Based on 99 replicates
Simulated p-value: 0.01
Alternative hypothesis: greater

     Std.Obs  Expectation     Variance
11.403084394  0.676852890  0.000281632
```

Site B 25 meters (hypolimion)

```
Monte-Carlo test
Call: RV.rtest(df1 = new_df, df2 = new_df2, nrepet = 99)

Observation: 0.9575236

Based on 99 replicates
Simulated p-value: 0.01
Alternative hypothesis: greater

     Std.Obs  Expectation     Variance
2.6858202391 0.8891274339 0.0006484995
```
